# Viral Burden: New insights into Estimating IgG Recognition Across Human Populations

**DOI:** 10.64898/2026.09.16.751894

**Authors:** Marek Adam Harhala, Katarzyna Gembara, Daniel C. Nelson, Andrzej Konieczny, Natalia Jędruchniewicz, Izabela Rybicka, Krystyna Dąbrowska

**Affiliations:** Institute of Human Biology and Evolution, Faculty of Biology, Adam Mickiewicz University, 61-615, Poznan, Poland; Laboratory of Phage Molecular Biology, Hirszfeld Institute of Immunology and Experimental Therapy, Polish Academy of Sciences, 53-114, Wroclaw, Poland; Research & Development Center, Regional Specialist Hospital, 53-114, Wroclaw, Poland; Institute for Bioscience and Biotechnology Research, University of Maryland, Rockville, 20850, MD, United States of America; Clinical Department of Nephrology, Transplantation Medicine and Internal Diseases, Institute of Internal Diseases, Wroclaw Medical University, 53-114, Wroclaw, Poland; Faculty of Medicine, Wroclaw University of Science and Technology, 53-114, Wroclaw, Poland

**Keywords:** Immunogenicity, Serological Profiles, Viral Antigens, VirScan, Population Immunity, Herd immunity, Vaccine

## Abstract

Assessment of viral burden in populations is essential for understanding virus epidemiology and herd immunity potential. While serological profiling is straightforward at the individual level, it remains challenging at scale. We propose analysis advancing in serological technologies for broader population-level analysis and comparison.

We used an epitope library in Phage Display ImmunoPrecipitation (PhIP) technology (VirScan type library) to assess IgG recognition of 49,630 representative viral oligopeptides in 134 serum samples from two populations (Poland and the US). Only 5.9% of oligopeptides were immunogenic, yet IgG recognition of viruses was consistent across populations—over 90% of virus species, 99% of genera, and 97% of families were detected. Shannon Diversity Index analysis supported these findings. Among immunogenic peptides, 9.1% were significantly more frequently recognized, though not correlated with recognition strength. We further proposed a normalization method to account for differences in viral proteome representation when assessing immune burden: the burden score. Finally, we demonstrated how this approach facilitates serological comparisons between populations.

These observations show that while people are exposed to similar viruses, they produce antibodies against different viral epitopes. This individual variability, combined with broad virus recognition, likely strengthens population-level protection. Epitope recognition frequency seems to be shaped more by population exposure than by magnitude of response that an epitope can induce. Accurately measuring viral burden can inform healthcare planning, and antiviral technologies development.

**Importance:** In this study, we explore how antibodies in human blood recognize representative proteome of viruses that attack humans, then characterize serological profiles in human populations. These profiles are crucial for modelling epidemiological factors and assessing viral burden within a population resulting in increased understanding of epidemiology in human populations. Presented results facilitate planning and predictions within medical service systems, including improvements in performance, capacity, and cost management. Moreover, a comprehensive understanding of virus burden can contribute to the development of more effective vaccines and therapeutics, guiding research towards areas of greatest need. Monitoring viral burden allows for the timely detection of emerging viral pathogens and their variants, enabling a rapid response to prevent outbreaks and minimize their impact on communities. We believe that leveraging Phage Display ImmunoPrecipitation technology used in our research may strongly support global efforts to control viruses in human populations.

## 1 Introduction

The human immune system exhibits a remarkable diversity in its responses to viral infections, mirroring the diversity among individuals themselves[1]. This biodiversity plays a pivotal role in shaping both the progression of viral infections and the corresponding immune reactions they elicit[2–4]. It significantly influences the range of interactions that determine the outcome of viral infections on a global scale, alongside temperature variations, other climate-related factors, and the presence of other pathogens[5,6]. Vaccination further contributes to this diversity. By stimulating the production of specific antibodies that can recognize viral particles and trigger immune responses, vaccines play a crucial role in protecting individuals from a wide range of infectious diseases[7]. This may complement, enhance, or even substitute for immunizations achieved after viral infections. Thanks to the somatic recombination of Ig-coding genes during B cell maturation, billions of antigens can be specifically recognised by IgG antibodies, which ultimately protect individuals from viral infections[8]. The combined impact of multiple factors contributes to the complexity of the clinical course of viral diseases, presenting a substantial challenge to epidemiological forecasting and modelling of viral transmission[9,10].

High-throughput identification of the complete array of antibodies circulating in human blood holds the promise of accurately predicting responses to specific viruses. It is difficult in an individual, but even more challenging at the societal level. A comprehensive understanding of immune phenotypes across entire populations could provide invaluable support for policymaking, epidemiological forecasting, healthcare burden management, and the planning healthcare expenditures within national budgets.

This holistic approach faces challenges due to the extreme variability of IgG antibodies in the human population. Traditional methods of IgG identification are often limited by the number of epitopes they can investigate, failing to fully capture the complexity of antiviral responses across populations. As a consequence, these methods may overlook potential correlations between responses to various viruses at both individual and societal levels. To address this gap, the Phage ImmunoPrecipitation Sequencing (PhIP-Seq) method has been developed and refined over the past decade[11,12]. This method led to the technology, which enables the detection and relative quantification of IgG fractions in human sera, with the capacity to identify tens of thousands of immunogenic viral epitopes[13,14]. At its core, this technology involves selecting short oligopeptides displayed on bacteriophage capsids (i.e., phage display), which interact with IgG antibodies present in serum samples. Oligopeptides that do not interact with IgG antibodies in the tested sera are subsequently removed, allowing for characterization of oligopeptides recognized by human IgG antibodies. In recent years, this technology has advanced rapidly, resulting in the development of dedicated software solutions, protocols, and significant review articles[15–17].

Exploring this technology, we aim to systematically assess its analytical capabilities through a systematic, data-driven approach. We characterize its immunomics potential (1) evaluating the serological landscape of the general population with respect to B-cell epitopes, (2) applying statistical methods to describe and interpret these profiles, (3) comparing patterns observed across distinct populations, and (4) examining the applicability of the viral burden concept within our dataset. Through these steps, we aim not only to explore the diversity and shared features of antiviral responses—across multiple biological scales, from oligopeptides to proteins, species, and families, but also to assess the practical utility of this technology for population-level immunomics.

## 2 Results

### 2.1 Large, non-random person-to-person diversity in specific recognition of viral oligopeptides

We evaluated the serological profile (IgG) in 134 serum samples from two test populations that are geographically separated: Poland (PL, 46 samples) and the United States (US, 88 samples). This was done by assessing the immunogenicity of 49,630 oligopeptide sequences, 56 amino acids in length, which represent the proteomes of 171 human-infecting viruses (a full list is available in Supplementary Data 1). A viral oligopeptide library was generated using phage display technology and immunoprecipitated with serum samples containing IgG antibodies. Specifically, IgG antibodies recognized the viral oligopeptides, forming IgG-oligopeptide complexes. Unbound oligopeptides were removed by washing. The isolated IgG-oligopeptide complexes, containing immunogenic oligopeptides, were analyzed to identify (count) how often each oligopeptide was recognized by specific IgG.

First, we assessed person-to-person diversity in the antiviral responses. An average of 90 ± 21 different immunogenic oligopeptides were recognized per serum sample (Figure 1A). Notably, the majority of tested oligopeptides showed no reactivity to the tested sera, with detected signals similar to or below those observed in the library before immunoprecipitation (i.e., input samples). Oligopeptides identified as immunogenic demonstrated signals above a cut-off value (adj. p < 0.05, false discovery rate FDR); of note, these signals showed a wide range of increased values, different between oligopeptides (Figure 1B).

**Figure 1.**
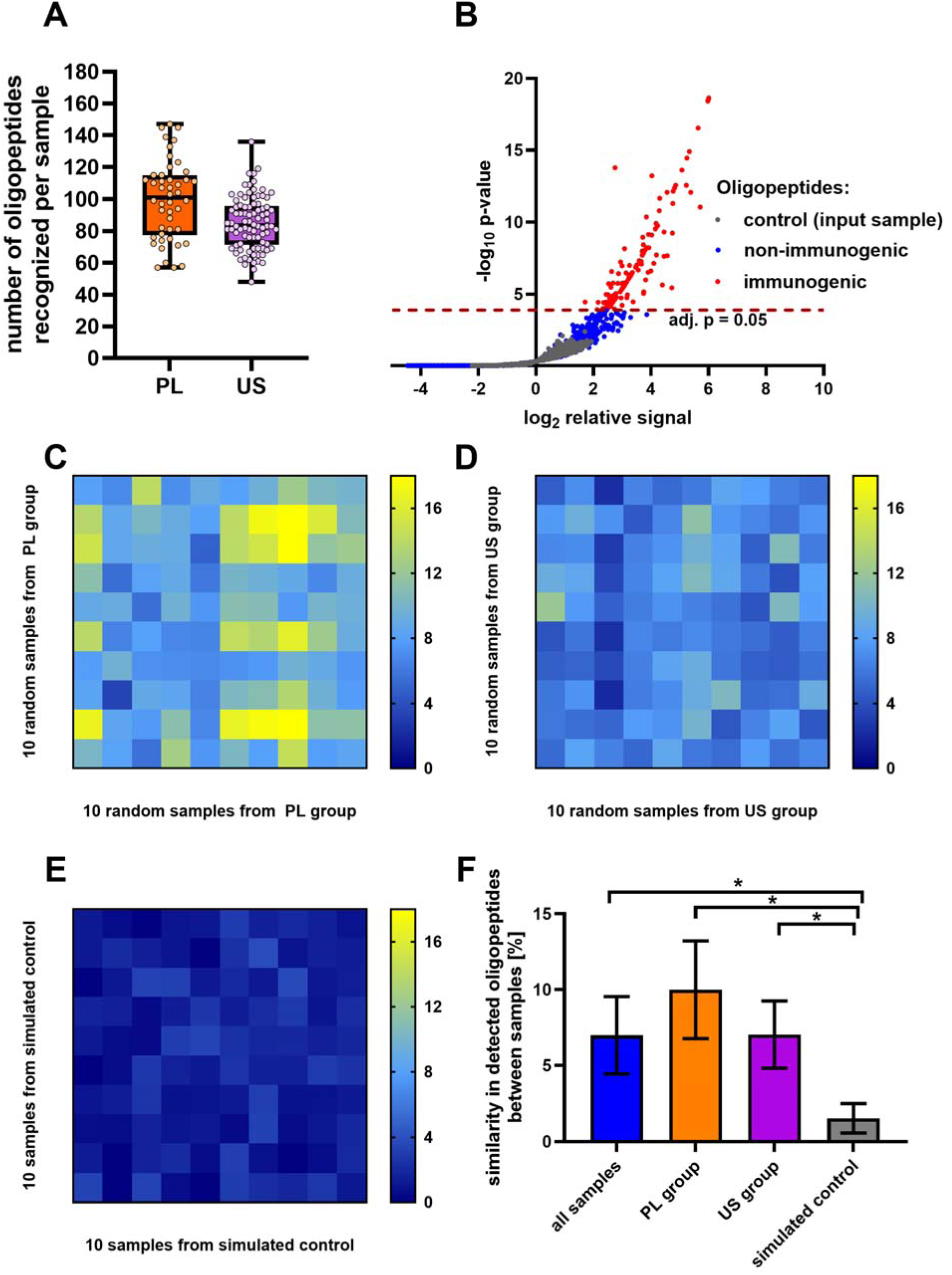
Characteristics of viral oligopeptides recognition in human sera from Poland (PL) and the United States (US). A viral proteome library consisting of 49,630 oligopeptide sequences, each 56 amino acids in length, was exposed to direct interaction with human IgGs. (A) Number of oligopeptides recognized by IgGs in the US (violet) and PL (orange) human sera, whiskers depict minimum and maximum number of detected immunogenic oligopeptides. Each dot represents one sample. (B) A volcano plot representing the typical serological profile (IgG) observed in this study. Each dot corresponds to a detected oligopeptide and is positioned to relative signal (enrichment) on the x-axis and the significance of this signal enrichment (p-value) on the y-axis. The relative signal is a fold change of a detected signal compared to the mean signal of all control samples (input samples, oligopeptide libraries before reaction with IgGs) for a given oligopeptide. P-values were calculated between the measured sample and all control samples (means and SD) for each oligopeptide separately. Grey dots represent oligopeptides detected in the control sample, red dots represent oligopeptides significantly enriched in the tested sample (immunogenic) (adj. p>0.05), blue dots represent oligopeptides not enriched in the tested sample (not immunogenic). Each dot represents a mean value of two technical replicates, with only oligopeptides significantly enriched in both replicates qualified as significantly enriched (immunogenic). (C) Heatmap presenting level of similarities between random samples in PL group, (D) the US group, (E) and in *in silico*-generated control. Color gradient corresponds to percentage of the same immunogenic oligopeptides detected between sample in a given row and column. (F) Level of similarity between samples in tested groups compared to an *in silico*-generated control. Bars and whiskers represent means and SD. Asterisk denote significant difference between groups, two-sided Welsh’s t test.

Our findings indicated that only 5.9% of the tested viral oligopeptides (2,926) were recognized as immunogenic, meaning they were recognized by human IgG in at least one of the serum samples (Supplementary Table 1 raw count data are in Supplementary Table 2). To put this in perspective, if oligopeptides in this experiment would be detected randomly, 21.7 ± 0.4% of these oligopeptides would be identified as immunogenic (see section 6.5 in Materials & Methods for details). This strongly supports specific selection of oligopeptides interacting with IgG (p<0.0001, binomial distribution, Supplementary Table 3). Nearly half of immunogenic viral oligopeptides (1,394, 47,6 %) were recognized by IgG in only one serum sample.

Comparisons between any two randomly selected samples revealed an average identity of recognized immunogenic oligopeptides as low as 6.5% ± 2.2%, approximately 6 oligopeptides out of 90; higher chance for identical recognitions was observed in PL samples (see also section 2.4 Geographical variations in serological profiles and virus recognition) (Figure 1C and 1D). This number is significantly higher than estimated for hypothetical random selection (1.5% ± 0.9%, p=4×10^-9^, see section 6.6 in Materials & Methods, Figure 1E). This suggests that, despite highly individualized serological profiles, human immune responses exhibit a minor but consistent preference for targeting certain viral epitopes (Figure 1F).

### 2.2 Recognition of virus species, genera, and families is largely shared across individuals, despite diverse individual responses to oligopeptides and proteins

We analyzed origins of each tested oligopeptide in terms of its viral protein, species, genus, and family (full list in Supplementary Table 4). If a tested oligopeptide was recognized as immunogenic in at least one serum sample, then the protein, species, genus, and family of origin for that viral oligopeptide were marked as “recognized” by IgG and “immunogenic” in that serum sample (Figure 2). Unlike individual oligopeptides, 94% of tested species (159 out of 170), 99% of genera (66 out of 67), and 97% of families (31 out of 32) were recognized (Figure 2A). However, at the protein level, a transient pattern was observed: over half of the tested proteins (1,667 out of 3,003), were non-reactive to IgGs in the investigated sera (Supplementary Tables 5-9). Thus, while only a small fraction (5.9%) of potential viral epitopes is recognized by human IgGs, this still allows for broad coverage of virus species, genera, and families by human immune responses.

**Figure 2.**
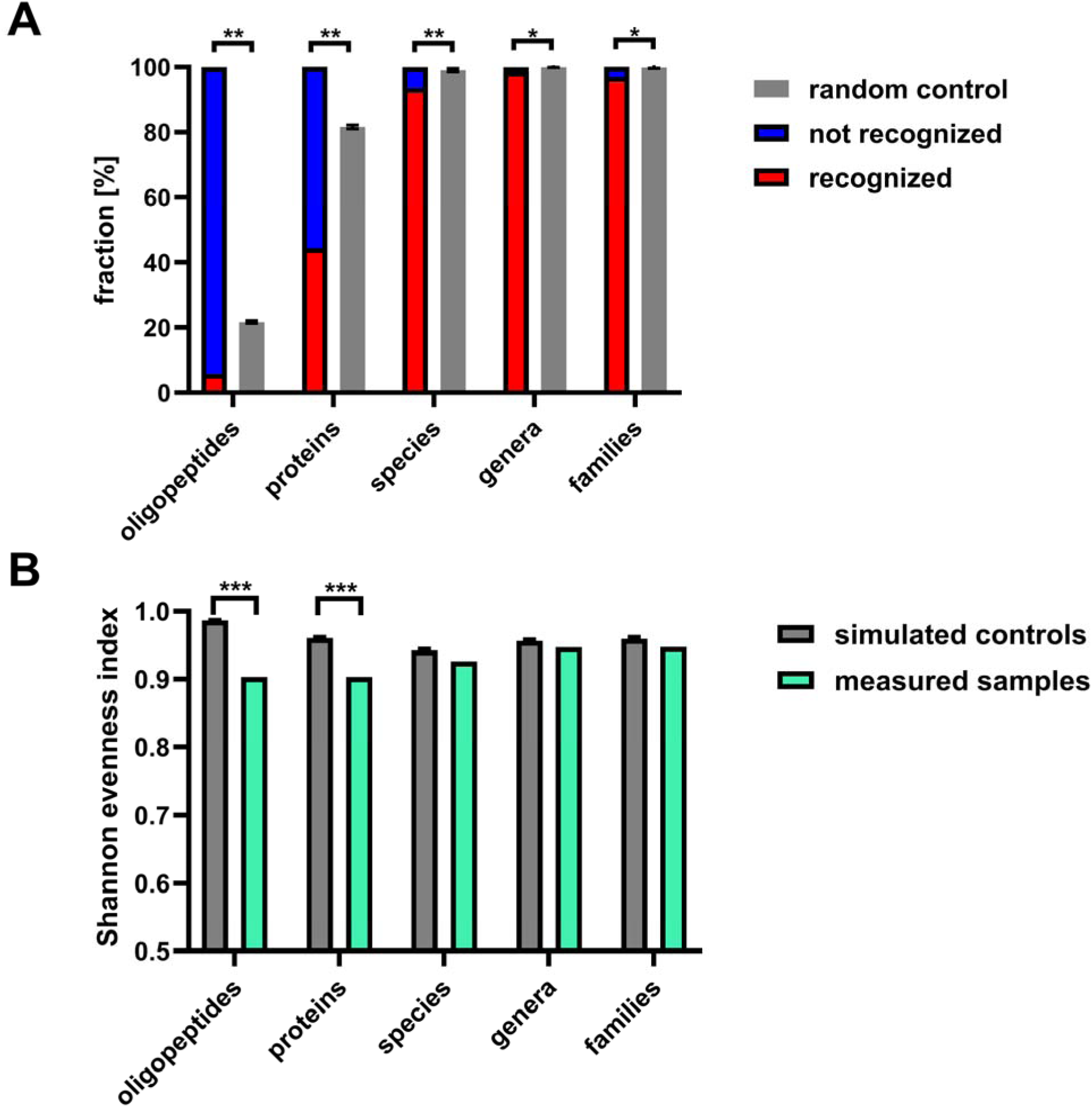
Taxonomic summary of viral oligopeptides found immunogenic in humans. (A) Fractions of viral oligopeptides and their corresponding proteins, species, genera, and families recognized by human IgG antibodies. Red bars represent the fractions of viral elements (oligopeptides and their parent proteins, species, genera, and families) that were recognized by IgG in at least one of the 134 human serum samples (immunogenic). Blue bars show the fractions not recognized in any tested sample. (B) Diversity in tested and simulated groups on all taxonomical levels, assessed with Shannon Evenness Index (SEI), that is Shannon Diversity Index (SDI) normalized to max SDI. Grey bars indicate expected fractions based on a model of random oligopeptide recognition (binomial distribution), serving as a null hypothesis. Whiskers represent standard deviations. Asterisks denote statistical significance of the differences between observed and expected recognition levels (** p□<□0.001, * p□<□0.05).

To assess the diversity within the serological profiles of the studied individuals, we used the Shannon Diversity Index (SDI) and its normalized value, SEI (Shannon evenness index, SDI divided by maximum SDI possible in tested sample). We compared the observed diversity with that expected under a random distribution (see section 6.8 in Materials & Methods). The diversity of serological profiles at the oligopeptide and protein levels was significantly lower than expected values computed by random distribution (Table 1, Figure 2B), indicating an uneven distribution characterized by the domination of certain oligopeptides or proteins (epitopes and antigens). However, at the levels of virus species, genera, and families, Shannon Diversity Index was indistinguishable from that expected by random distribution (Table 1, Figure 2B). Thus, despite substantial person-to-person diversity in specific recognition of viral oligopeptides or proteins, the overall recognition of a vast majority of virus species, genera, and families appears unified at the population level. This suggests that while different human individuals respond to the same (or similar) viruses, they do so with antibodies targeting different epitopes and, to some extent, different antigens, within viral particles.

**Table 1:**
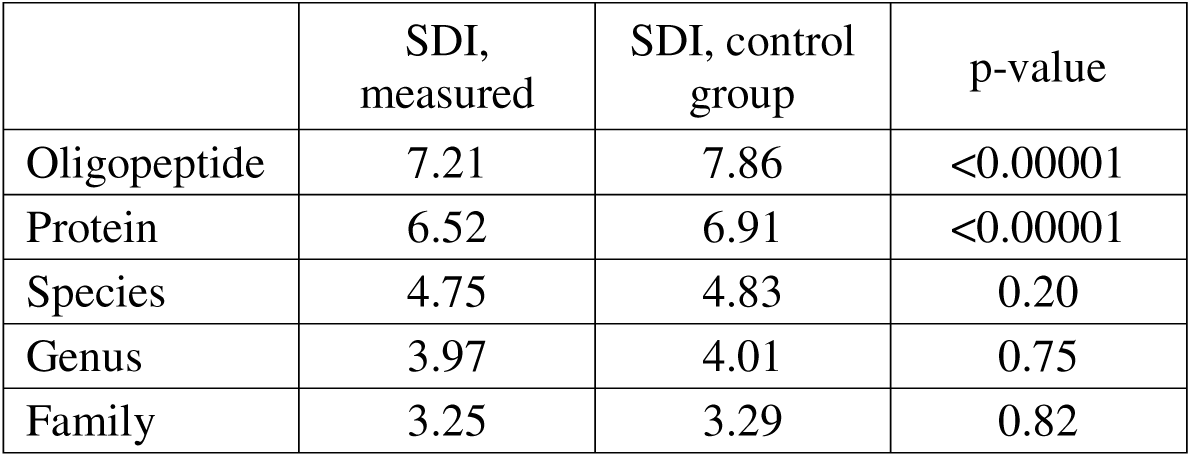
Comparison of the Shannon Diversity Index (SDI) for viral immunogenic oligopeptides. A comparison of the SDI between the investigated population and expected values from a computed control assuming random distribution of immunogenic oligopeptides in the tested samples. Comparisons are presented for viral immunogenic oligopeptides and their corresponding proteins, species, genera, and families of origin. The “SDI, measured” column represents SDI calculated for the tested population, while the “SDI, control group” column represents expected SDI values. P-values are calculated to determine the significance of differences between observed values and the expected values from the computed random distribution control (see section 6.8 in Materials & Methods).

It should be emphasized that this analysis was performed directly, with the basic analytical unit defined as a single epitope and its association with proteins, species, and higher taxonomic levels. In the further part of the study (sections 2.5 and 2.6), the analysis was extended to identify individual viral species that likely represent true exposure events and the corresponding induction of antibodies by these viruses.

### 2.3 Magnitude of response has a limited effect on how often an epitope is recognized across the population

Unlike the majority of immunogenic oligopeptides, some are commonly recognized by IgG across multiple samples (Figure 3A). To characterize these frequently recognized oligopeptides, we calculated the average probability of IgG recognition across serum samples. From this analysis, we identified oligopeptides that were over- or underrepresented. Only 265 oligopeptides (9.1% of all immunogenic oligopeptides) were significantly overrepresented in the recognized fraction, while none were underrepresented (Figure 3B, adjusted p-value<0.05, false discovery rate, see section 6.9 in Materials & Methods for details).

**Figure 3.**
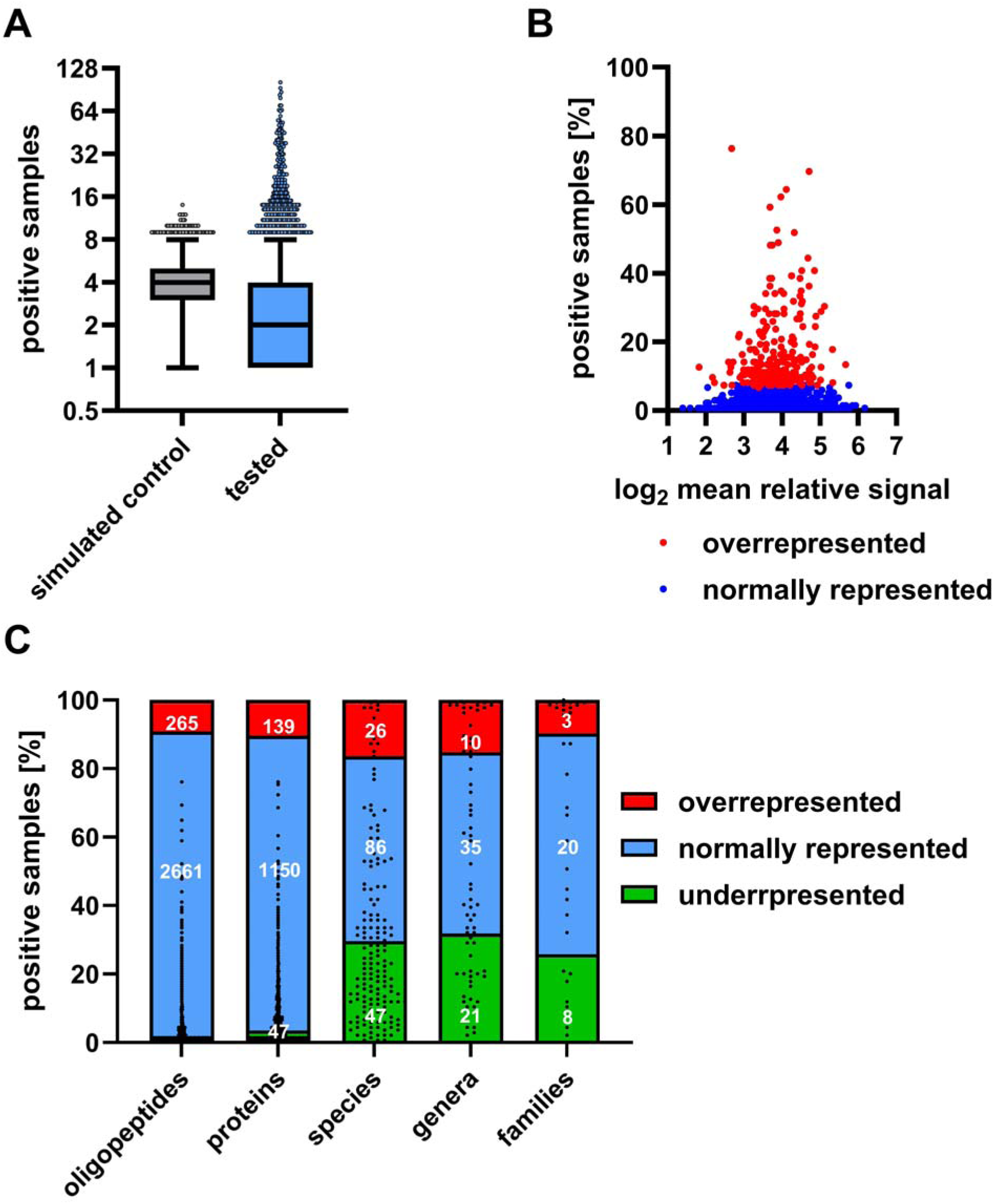
Relation between immunogenicity (relative signal) and recognition frequency of viral oligopeptides. Analysis of the recognition frequency and immunogenicity (relative signal) of the investigated oligopeptides was used to identify immunogenic oligopeptides that were overrepresented in the tested population. (A) Number of samples in which an immunogenic oligopeptide was detected. Simulated control (grey) and tested group (blue) are presented. Boxplot and whiskers show means and 99% interval of all 2,969 detected immunogenic oligopeptides. Dots represent top 1% of most often recognized immunogenic oligopeptides. (B) Relation between number of samples in which immunogenic oligopeptides were detected and relative signal of these oligopeptides. Each dot represents an immunogenic oligopeptide, with those significantly overrepresented (‘*overrecognized’*) in the population shown in red (adjusted p<0.05) and others shown in blue. (C) Summary of overrepresentation analysis. Stacked boxplot shows percentages of immunogenic oligopeptides, proteins, species, genera and families over or underrepresented in our tested sera. White numbers indicate counts of viral elements under-, over- or normally represented. Each black dot represents a viral element and shows percentage of samples that recognized this individual viral element. If the oligopeptide was immunogenic, its protein, species, genera and family of origin was also marked as immunogenic. The cut-off value (adjusted p<0.05) for over- and underrepresented entities was calculated by comparing the observed recognition frequency to the theoretical frequency expected under the assumption of random recognition of immunogenic oligopeptides, proteins, species genera and families in tested samples.

Importantly, overrepresentation of immunogenic oligopeptides did not correspond to the strongest average immunogenic response. (Figure 3B). We observed very weak correlation between the frequency of recognition and the magnitude of response, defined as the mean increase of relative signal (two tailed correlation coefficient r(2,924) = 0.19, 95% CI 0.15 - 0.22; Spearman’s ρ(2,924) = 0.34, p<0.0001). This suggests that viruses recognized in a large fraction of the population do not induce very strong response on average. Viruses with the strongest immunogenic epitopes capable for high magnitude of response are not the most commonly recognized in the tested population. Notably, in some samples these overrepresented oligopeptides elicited a relatively strong response, but even then, overrepresented oligopeptides demonstrate overall medium increase in the relative signal (Supplementary Data 10). Recognized viral proteins, species, genera, and families showed a more complex pattern of recognition. While some were overrepresented, an increasing number also appeared underrepresented (Table 2, Figure 3C). This corresponds to differences in viral epidemiology and possibly differentiated viral strategies of immune evasion as primary factors contributing to the limited recognition of viral groups in human populations.

**Table 2:**
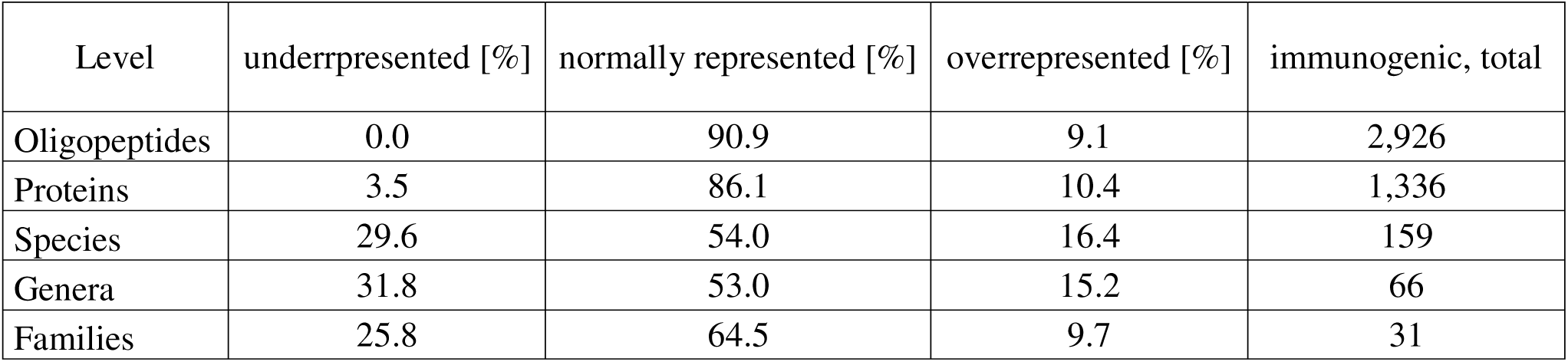
Frequency distribution of immunogenic viral components. Distribution of immunogenic viral oligopeptides, proteins, species, genera, and families that show significantly overrepresented or underrepresented frequencies within tested population. These fractions were identified based on statistically significant differences between observed and expected frequencies, with the latter calculated under the assumption of random recognition of immunogenic oligopeptides in the population (see section 6.9 in Materials & Methods).

| Level | underrpresented [%] | normally represented [%] | overrepresented [%] | immunogenic, total |
| --- | --- | --- | --- | --- |
| Oligopeptides | 0.0 | 90.9 | 9.1 | 2,926 |
| Proteins | 3.5 | 86.1 | 10.4 | 1,336 |
| Species | 29.6 | 54.0 | 16.4 | 159 |
| Genera | 31.8 | 53.0 | 15.2 | 66 |
| Families | 25.8 | 64.5 | 9.7 | 31 |

### 2.4 Geographical variations in serological profiles and virus recognition

The tested serum samples were then analyzed as two separate groups: one from the United States (US, n=88) and one from Poland (PL, n=46) (demographic information on the participants is available in Supplementary Table 11). Our analysis of viral oligopeptide recognition shows how serological profiles may differ between distinct populations. The analysis revealed that 32% of the immunogenic oligopeptides were recognized by IgG in both groups (938 out of 2,926). However, 47% (1,361 out of 2,926) were recognized only by the US group, and 21% (627 out of 2,926) only by the Polish group (Figure 4A, Table 3).

**Figure 4.**
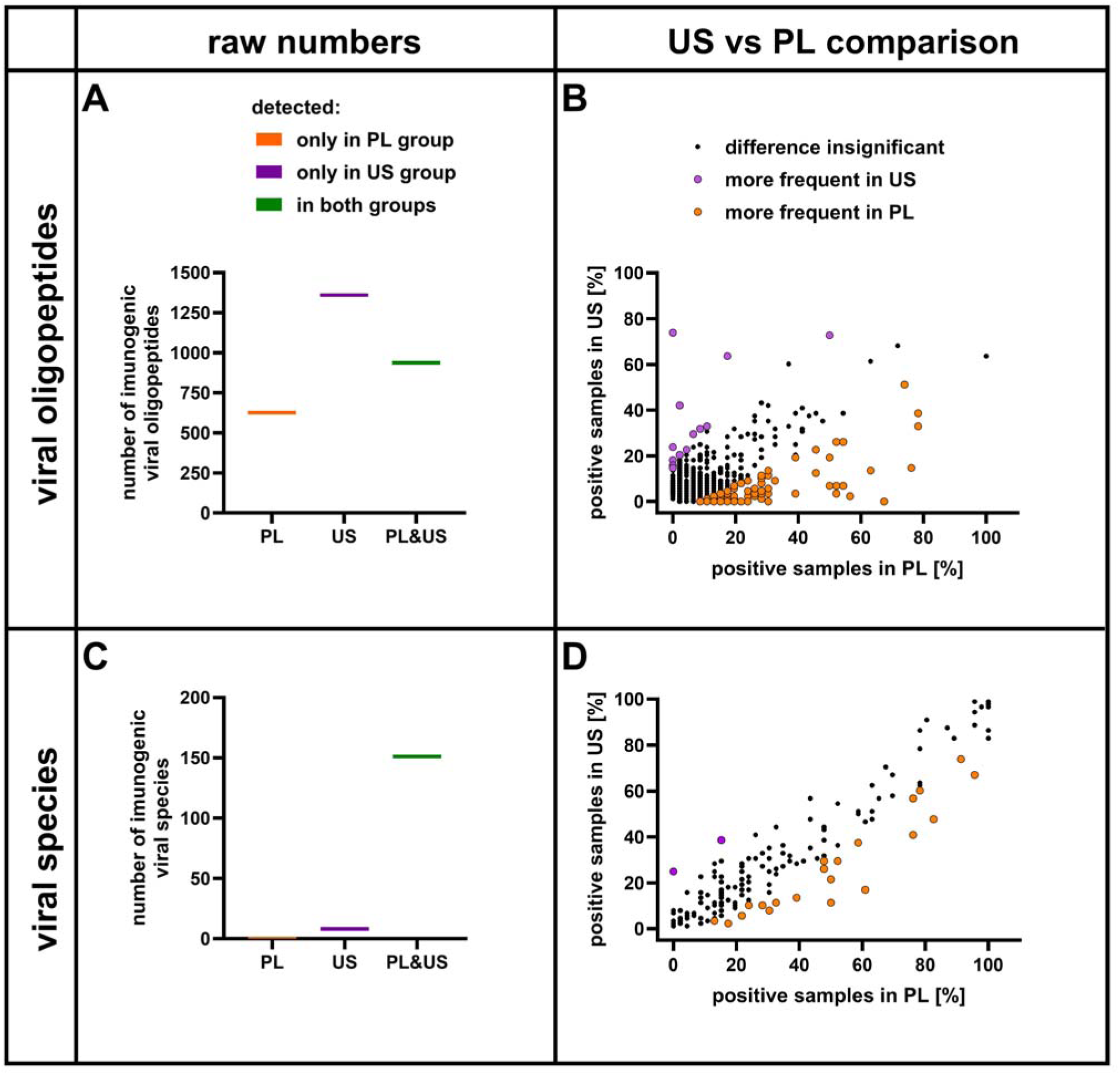
Comparative overview of immunogenicity metrics in geographically distinct populations. (A) Numbers of oligopeptides recognized by human IgGs (immunogenic). (B) Comparative detection plot for immunogenic oligopeptides in PL group (x-axis) and in the US (y-axis) group. (C) Numbers of viral species recognized by human IgG. (D) Comparative detection plot of viral species associated with immunogenic oligopeptides in PL (x-axis) and in the US (y-axis) group. Violet and orange dots show oligopeptides or virus species recognized significantly more often in the US or PL group, respectively (adj. p-value < 0.05). Adjusted p-value was calculated by comparing probabilities of measured frequencies to mean and SD of all detected immunogenic oligopeptides and species.

**Table 3:** Comparison of regional serological profiles between populations of Poland (PL) and the United States (US), oligopeptides. The table presents the number of immunogenic oligopeptides recognized by IgG in human sera, categorized as follows: recognized only in Polish donors (“PL only”), only in US donors (“US only”), or in both groups (“both groups”). These numbers are further divided into three categories based on statistical analysis to identify differences between the geographically distinct groups. The rows labeled “more frequent in US, adj. p<0.05” and “more frequent in PL, adj. p<0.05” represent number of immunogenic oligopeptides that were significantly more abundant in the US or PL group, respectively, and rarely or not at all recognized in the other group. The row labeled “difference non-significant, adj. p > 0.05” shows the number of immunogenic oligopeptides with no significant differences between groups, which may be attributed to large variances between individuals. The “total” column/row represents all detected immunogenic oligopeptides in a respective row/column.

| statistical significance of differences between US and PL group | number of oligopeptides recognized by IgG in: |  |  |  |
| --- | --- | --- | --- | --- |
|  | both groups | US only | PL only | total |
| difference non-significant, adj. $p > 0.05$ | 866 | 1,354 | 594 | **2,814** |
| more frequent in US, adj. $p < 0.05$ | 8 | 7 | 0 | **15** |
| more frequent in PL, adj. $p < 0.05$ | 64 | 0 | 33 | **97** |
| **total** | **938** | **1,361** | **627** | **2,926** |

Some of the observed differences could have arisen by chance because the serum samples represented only subsets of two large and heterogeneous national populations. To account for this sampling variability, we used a binomial model to estimate the probability that the differences in recognition frequency between the US and Polish groups occurred by chance (see Materials and Methods, Section 6.10). This analysis identified 112 immunogenic oligopeptides (3.8% of all immunogenic oligopeptides) that were recognized significantly more frequently in one group than in the other (adjusted p < 0.05). (Figure 4B, Table 3, Supplementary Table 12, see Section 6.10 in Materials & Methods). Notably, 97 of these 112 oligopeptides were overrepresented in the Polish population only (Table 3).

On average, an average immunogenic oligopeptide was recognized in 3.7% of serum samples in the US group and in 6.4% of the Polish group (p<2×10^-16^, Wilcoxon rank sum test). We hypothesize this being a consequence of Polish population being very homogeneous in terms of ethnic origin (97% Western Slavic, white)[18] and Poland being much smaller than the US (3.2% of the US land area). A group from United States is far more diverse ethnically than PL (for details see Supplementary Table 11). This supports the hypothesis that more diverse populations also show greater diversity in IgG profiles. Although both countries are classified by the World Health Organization as high-income, socioeconomic and other factors likely contribute to the observed differences[19,20].

We observed small but significant differences in the frequency of virus species recognition between tested populations (Supplementary Table 13). Of the 159 identified virus species, 95% (151 species) were recognized in both groups, while the remaining 8 species were recognized only in the US group (Figure 4C, Table 3). However, only 14 % of all identified virus species (23 out of 159) were recognized more frequently in either the US (2 species) or Polish group (21 species) (Figure 4D, Table 4). Interestingly, the only virus species that was uniquely recognized in the US group and in a sufficient number of samples to make the difference statistically significant (p□<□0.05) was Human/Bovine parainfluenza 3 virus (Supplementary Table 13). Two oligopeptides derived from the same viral fusion glycoprotein (UniRef90_P06828) are the source of this observed US vs. PL difference. One is more frequently recognized and stronger (UniRef90_P06828_oligopeptide_19, detected in 21 samples, relative signal between 26 and 6.8) and the other is weaker (UniRef90_P06828_oligopeptide_4, detected in 3 samples, relative signal between 7.77 and 6.2). In total, 31 oligopeptides representing this virus were tested and only two were found immunogenic. Noteworthy, the stronger one is a terminal oligopeptide of their protein of origin. We propose that this observation is consistent with the US being the largest producer and exporter of beef products in the world, with a per capita consumption of over 26 kg of beef products in 2023, compared to only 1.7 kg per capita in Poland. We speculate that in the US population may be exposed more often to such viral material, likely as a non-infectious antigen present in the environment, for example through food or meat processing. This suggests that differences in immune responses due to varying levels of exposure to viruses can be detected using the analysis pipeline proposed here.

**Table 4:** Comparison of regional serological profiles between populations of Poland (PL) and the United States (US), species. The table presents numbers of virus species recognized by IgG in human sera, categorized as follows: recognized only in Polish donors (“PL only”), only in US donors (“US only”), or in both groups (“both groups”). These numbers are further divided into three categories based on statistical analysis to assess differences between geographically distinct groups. The rows labeled “more frequent in US, adj. p<0.05” and “more frequent in PL, adj. p<0.05” indicate the number of virus species that were significantly more abundant in the US or PL group, respectively, and rarely or not at all recognized in the other group. The row labeled “difference non-significant, adj. p > 0.05” shows the number of virus species with no significant differences between the groups, which may be attributed to large variances between individuals. The “total” column/row represents all detected virus species in each respective row/column.

| statistical significance of difference in frequency between US and PL group | species recognized by IgG in: |  |  |  |
| --- | --- | --- | --- | --- |
|  | both groups | US only | PL only | total |
| difference non-significant, adj. $p > 0.05$ | 129 | 7 | 0 | **136** |
| more frequent in US, adj. $p < 0.05$ | 1 | 1 | 0 | **2** |
| more frequent in PL, adj. $p < 0.05$ | 21 | 0 | 0 | **21** |
| **total** | **151** | **8** | **0** | **159** |

### 2.5 Normalization strategy for assessing viral burden on the immune system

We identified viruses that exert the most significant pressure on immune systems in the studied population following the methodology of Xu et al.[13,14]. Briefly, a virus-specific threshold that accommodated differences in viral proteomes size was calculated, establishing a cut-off number of recognized epitopes (per sample) indicating exposure of the serum donor to this virus species. This assumption was made to reduce the influence of cross-reactive antibodies, which can recognize viral epitopes even when induced by different antigens. Such an approach is relevant for both symptomatic and asymptomatic infections, as well as vaccinations.

We identified 70 virus species (41.2% of all 170 tested virus species) for which exposure defined as by Xu et al. was indicated in at least one serum sample[13,14]. These viruses were represented by 1516 immunogenic oligopeptides (52% of all 2,926 immunogenic oligopeptides detected in the presented study). We observed marked differences in the distribution of IgG recognition across the viruses. For example, responses to the seven most common virus species were detected in at least 40% of individuals, while another 15 were detected in at least 10%. Out of the 70 identified virus species, responses to 22 (31.4%) were found in only one individual (Supplementary Table 14).

Importantly, we observed substantial variation in how different viruses are represented in the databases. Some virus species are highly represented with many proteomic records, while others are not. We observed that the frequency of detected recognition correlated with the number of oligopeptides available in databases for a given virus. For instance, seven most commonly recognized virus species represented 30.1% of all immunogenic oligopeptides detected in any tested serum samples. Statistical analysis showed that the number of serum samples showing exposition to a virus was positively correlated with the number of oligopeptides representing this specific virus in the original library (Figure 5A, Pearson r(68)=0.73, 95% CI: from 0.59 to 0.82, Spearman’s ρ(68) = 0.69, p-value<0.0001). Thus, assessing viral burden simply by calculating fraction of serum samples recognizing a virus was insufficient. As an example, Human Immunodeficiency Virus 1 (HIV-1) is represented by 2,743 oligopeptides, accounting for over 5% of all oligopeptides and making it the most widely represented virus species. When calculated without correction, it appeared to be one of the high-burden viruses, with 32% of tested samples showing a positive response. This finding is inconsistent with epidemiological data from the World Health Organization, which estimates the prevalence to be approximately 0.5%[21].

**Figure 5.**
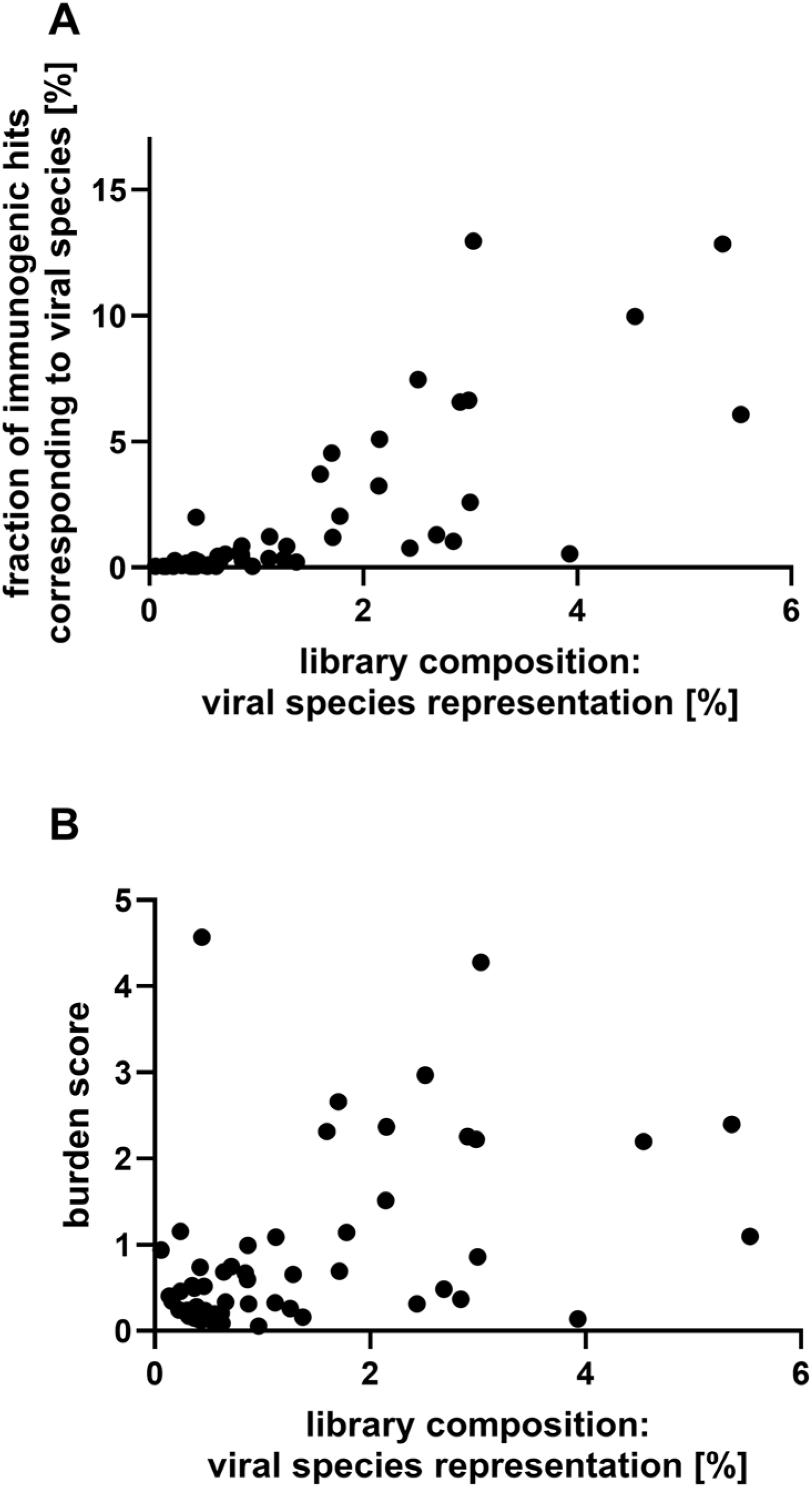
Impact of input oligopeptide representation in tested library on virus-specific detection rates. (A) Calculation without normalization to initial library composition. (B) Calculation normalized to initial library composition (the burden score) Each dot represent virus species to which at least one serum sample was exposed according to methodology by Xu et al.[13,14]

To address this issue, for evaluating the viral burden on immune systems using epitope libraries in Phage Display ImmunoPrecipitation (PhIP) technologies, we propose extending the analysis with a calculation of the **burden score**. High burden score indicates virus species that are recognized widely in the tested population. High-burden is defined in this study as significantly higher burden score. We define burden score as the ratio of two fractions: the fraction of detected immunogenic oligopeptides representing a specific virus species, divided by the fraction of oligopeptides representing this specific virus species in the original library (see details in section 6.11 of Materials & Methods). This correction serves as data normalization. With this applied correction, we present a list of high-burden virus species identified in the investigated populations (full list in Supplementary Table 14). We evaluated the viral burden score results using the same approach as described in the previous paragraph. The correlation of burden score with the number of oligopeptides representing this specific virus in the original library (Figure 5B, Pearson r(68)=-0.26, 95% CI: from −0.47.to −0.03, p-value=0.029, Spearman’s ρ(68) = −0.37, p-value=0.0018) is far lower than in the analysis where the burden score was not applied (p<0.0001, Fisher r-to-z transformation). This suggests that proposed burden score provides an improved assessment of viral burden than for the previously used number of serum samples exposed to a virus species. Also, Human Immunodeficiency Virus 1, despite being most represented virus species in the original library, showed no significant increase in burden score with the proposed calculations as expected based on WHO estimated prevalence.

Notably, the burden score shows significant results for underrepresented viruses. For example, human respiratory syncytial virus is represented by only 217 oligopeptides in the original library (0.44% of all tested, 46^th^ rank) with only 19 of them recognized by IgG in this study as immunogenic oligopeptides. Despite this, 22% of serum samples showed exposure to human respiratory syncytial virus (14^th^ rank) and it received the highest burden score in our analysis.

### 2.6 High viral burden is correlated with the recognition frequency of immunogenic epitopes, but not with their strength

For the purposes of this study, we define epitope strength as its capacity to induce higher (strong epitope) or lower (weak epitope) magnitude of specific IgG levels in an individual. Recognition frequency represents how often an epitope has been recognized in a population. We assessed potential correlation between the average strength of response to recognized oligopeptides and the frequency of recognition of relevant virus species based on the burden score and we found no significant correlation (Pearson’s r(2,924)= −0.06, 95% CI −0.30 to 0.19, p=0.64, spearman’s ρ(2,924) =0.11, p=0.41).

However, we observed a significantly higher frequency of immunogenic oligopeptides across the proteomes of high-burden virus species. The fraction of recognized oligopeptides in the proteomes of high-burden viruses (11 virus species) was 6.7% ± 1.5% on average. This was significantly higher than in other virus species recognized in more than 5% of serum samples, but with lower or moderate burden score (15 virus species), where the average fraction was 3.35% ± 0.92% recognized oligopeptides. This difference was statistically significant (p-value = 8.7 × 10□□, two-tailed Mann–Whitney test; see Figure 6). We conclude that the high-burden virus species are not those enriched in strong epitopes, but rather those characterized by a larger number of epitopes that can be recognized in many individuals.

**Figure 6.**
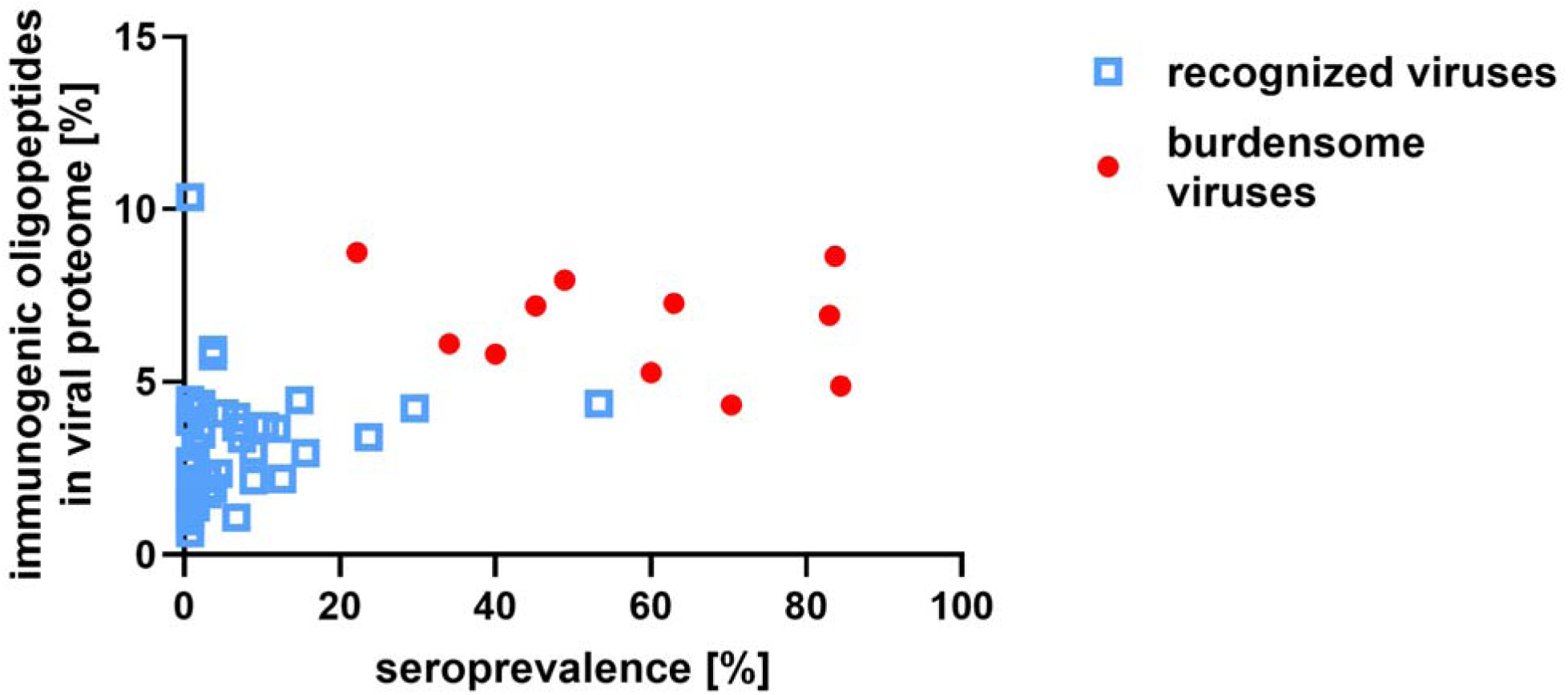
Immunogenicity of viral proteomes vs. population-level seroprevalence. Seroprevalence-fraction of serum samples with IgG recognition of ≥3 oligopeptides from a given virus, proteome immunogenicity-fraction of immunogenic peptides among all tested peptides from the virus. Each dot represents a virus species, red dot – burdensome virus species, blue square – non-burdensome but recognized virus species.

## 3 Discussion

In this study, we explored viral epitope library in PhIP technology as a tool to understand and characterize serological profiles in human populations. These profiles serve as valuable source of data for elucidating crucial epidemiological factors, such as population or herd immunity, and for assessing viral burden within a population. To place the presented results in the proper context, it is important to acknowledge the specific limitations of this method. This technology relies on libraries of short, linear peptides, which limits the detection of antibodies that recognize conformational or discontinuous epitopes, as well as those dependent on post-translational modifications such as glycosylation. In addition, incomplete or uneven representation of viral proteomes in the library, together with variability in phage display efficiency, may introduce biases in epitope coverage and detection sensitivity. Sequence similarity among related viruses can further lead to antibody cross-reactivity, making it difficult to assign signals to specific pathogens. Moreover, the readout is based on enrichment and sequencing, providing relative rather than absolute measures of antibody binding, and does not directly reflect antibody affinity, concentration, or neutralizing activity. Also, at the population level, mean relative signals may obscure interindividual heterogeneity and may not fully capture strong responses limited to individual samples. Finally, the method captures systemic humoral responses but does not account for cellular immunity or tissue-specific antibody repertoires. Despite these limitations, the approach offers substantial potential to advance our understanding of the molecular interactions underlying population-level immune responses, providing a powerful framework for population-scale immunological discovery[15–17,22–24].

Our findings show that only 5.9% of the tested viral oligopeptides (2,926 in total) were recognized by human IgG in any of the investigated serum samples. Nearly half of these (1,394 oligopeptides) were recognized in only one serum sample. This suggests that most individual serological profiles are unique and that humans (as a population) respond to a wider range of viral oligopeptides, but with only a small subset of "universal oligopeptides" recognized across multiple individuals (Supplementary Table 5, Figure 1). This is consistent with previous studies using the same technology, which demonstrated substantial differences in epitopes recognized by anti–Epstein–Barr virus (EBV) responders, in the recognition of seasonal coronaviruses (HCoVs), and in responses to other viruses[22–24].

However, when we analyzed the origin of each tested oligopeptide—by protein, species, genus, and family—we observed broad and increasing coverage of recognized proteins and taxa across the tested population. (Supplementary Tables 6-9). In contrast to oligopeptides, over 90% of the tested species, genera, and families were recognized (Figure 2). This finding is further supported by calculations of the Shannon Diversity Index, which showed a statistically significant decrease in the diversity of recognized viral oligopeptides compared to expected values from a simulated random distribution. At the same time, diversity at the level of virus species, genera, and families remained unchanged (Table 1). Of note, SDI values are computed using random controls derived from properties of the observed data, including the number of detected oligopeptides per sample and associated p-values, and therefore already account for differences in sequencing depth. In addition, we minimized sequencing-depth bias by using signal-based measures instead of raw counts and by comparing post-immunoprecipitation data to the input library for p-value estimation, rather than relying on sequencing-depth-dependent models.

To properly address the observation of relatively high coverage of viral species, genera, and families observed in this study, we acknowledge that antibody binding signals detected by PhIP-based approaches should be interpreted with caution. Such assays measure binding rather than functional activity, and therefore may capture a mixture of biologically relevant and non-relevant interactions. In particular, part of the observed reactivity may reflect cross-reactive antibodies elicited by unrelated antigenic exposures, as well as background binding that does not translate into neutralization or protective immunity. At the same time, we cannot exclude that a subset of these signals represents biologically meaningful responses, including potentially natural antibodies. Natural antibodies are predominantly of the IgM isotype, but they may include IgG isotype[25]. Further, immunoprecipitation using Protein A or Protein G shows a strong preference for IgG, though IgM binding is not entirely excluded[26]. Accordingly, we consider that the signal detected by PhIP likely comprises a combination cross-reactivity of uncertain relevance, and natural antibody responses, the functional significance of which remains to be determined in future studies.

Despite significant molecular variability in immune responses from person to person, the immune responses to virus species and higher taxonomic groups appear more unified at the population level. This implies that while different individuals may respond to the same or similar viruses, their antibodies target distinct epitopes within viral particles. This variability arises from natural differences between individuals, including HLA diversity, antigen processing pathways, and immune history, all of which shape epitope recognition patterns[27]. The concept of immunodominance hierarchies indicates that the immune system does not respond equally to all fragments (epitopes) of an antigen. Instead, some epitopes are more efficient in inducing specific response that others. Our results indicate that the pattern of epitope targeting is highly individualized. Rather than contradicting the concept of immunodominance, these observations support its substantial variability across individuals. Such diversity is also important for addressing the natural variability in individual immune responses and for reducing the risk of viral escape from vaccine-induced herd immunity. A broader repertoire of viral epitopes seems to increases the likelihood of eliciting effective immune responses across diverse populations, although further research is needed to explore this concept in detail.

We observed that certain viral oligopeptides, proteins, and taxonomic groups (species, genera, and families) were more frequently recognized by IgG antibodies than others. To explore this further, we calculated the average probability of each viral component being recognized by IgG in the tested serum samples and compared these probabilities to the actual frequencies of recognition across the population. This analysis showed that only 9.1% of all immunogenic oligopeptides were significantly overrepresented ("overrecognized") in the studied population, while none were significantly underrepresented. Most virus species and higher taxonomic groups showed no statistically significant over- or underrepresentation, accounting for nearly 60% of the total. However, we noted a gradual increase in the fraction of underrepresented groups at higher taxonomic levels (Figure 3).

Interestingly, among the identified immunogenic oligopeptides, we found only a weak correlation between the frequency of recognition and the mean relative signal intensity (representing the magnitude/strength of response). This finding suggests that the magnitude of response that viral epitopes are able to induce has a limited influence on how frequently they are recognized in the population (Figure 3) [28,30]. It further implies that additional biological and medical factors, beyond physicochemical or other properties of specific epitopes, efficiently shape epitope recognition patterns in humans [30].

The epitope-level analysis was extended to identify viral species that likely reflect true exposure and the corresponding induction of antibodies. This enables the identification of high-burden viruses, defined as those recognized significantly more frequently in the population than others. We followed the approach described by Xu et al. [13,14]. Six viruses, comprising just 3.5% of all virus species analyzed, made an overwhelming contribution to the group most frequently recognized in the studied population (Supplementary Table 14). These same six viruses also accounted for 28.2% of all immunogenic oligopeptides detected in the tested serum samples. Importantly, we also noted considerable variation in how different viruses were represented in the reference databases. For example, the ten most represented virus species accounted for 37% of all tested oligopeptides, while the ten least represented species made up only 0.3%. As a result, the frequency of viral recognition was strongly correlated with the number of oligopeptides available for each virus. Other manuscripts struggled with the same inequalities in virus representation however, not all publish data sufficient for such evaluations. Monaco et al., 2022, shows that some virus species can be represented by up 5% of oligopeptides while other virus species are represented by less than 0.001% of olidopeptides [17]. The manuscript we based our work also show similar inequality due to use of similar database [14]. To account for this bias and better estimate viral burden, we introduced a **burden score**. This score represents the prevalence of immune recognition of a given viral species relative to other viruses and to the total number of oligopeptides assigned to that virus in the library. Applying the burden score allowed us to more accurately identify the viruses most strongly recognized in the study population (Table 5).

**Table 5:**
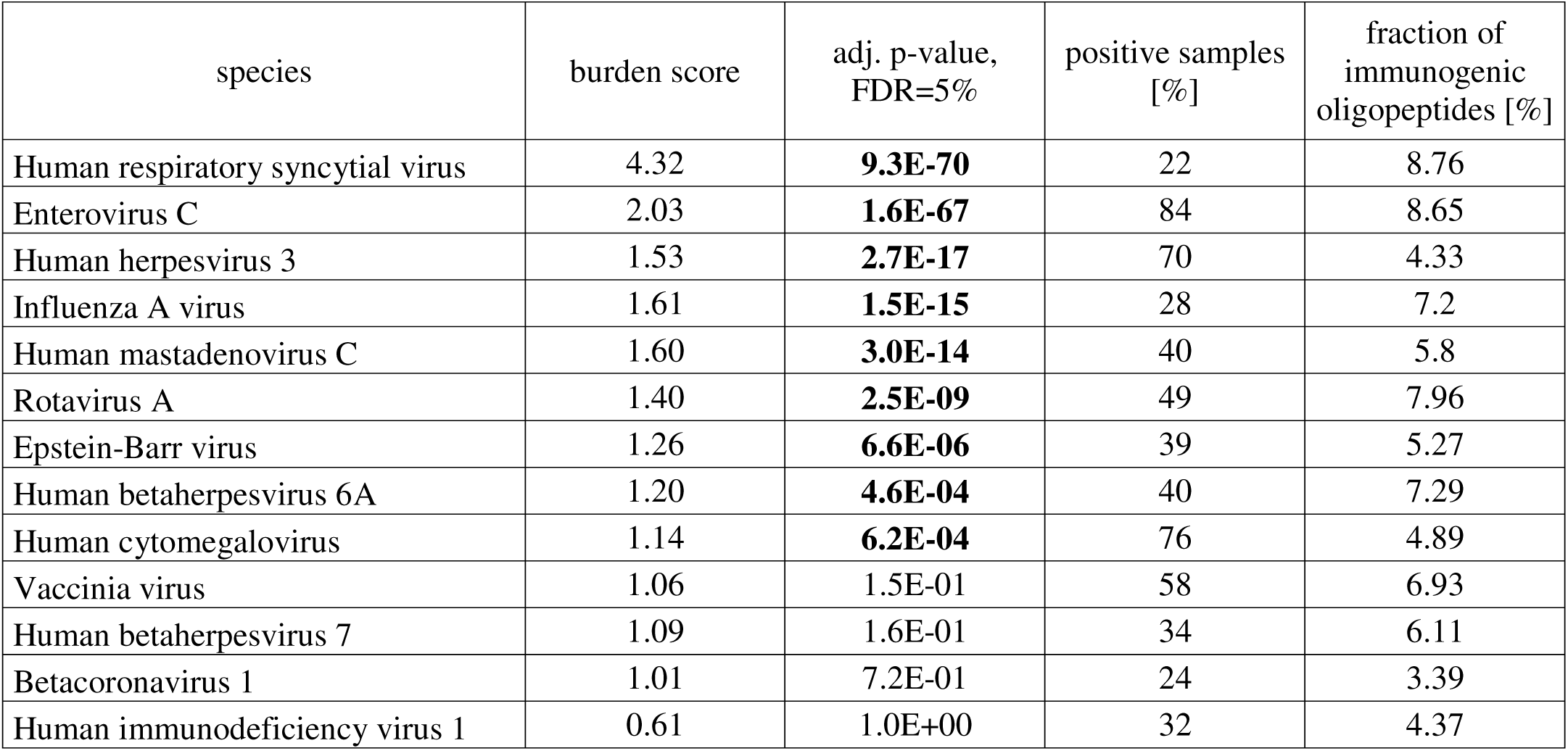
Impact of common virus species on the human population. This table provides an overview of virus species in the tested population, along with an analysis of the burden score to account for accidental cross-reactions due to the quantity of oligopeptides in the tested library. The “burden score” column indicates the odds of recognizing oligopeptides representing specific virus species as immunogenic compared to finding them in the original library by chance. A burden score significantly greater than one suggests a high viral burden with a low likelihood of accidental cross-reactions. The “positive samples [%]” column shows the percentage of tested sera that reacted to three or more oligopeptides of a specified virus species, following the method described by Xu et al., 2015[14]. Only virus species recognized in more than 20% of tested sera are listed. The “adj. p-value” column represents the probability that a burden score of the observed value or greater occurred by chance, using a one-tailed binomial distribution. Values where p<0.05 are highlighted in bold. Finally, the “fraction of immunogenic oligopeptides [%]” column indicates the percentage of oligopeptides representing specific virus species that were identified as immunogenic.

We believe that accurately assessing virus burden across populations provides information crucial for understanding the epidemiology of different viruses and herd immunity in humans. This can facilitate planning and predictions within medical service systems, including improvements in performance, capacity, and cost management. Such insights can inform public health interventions, aiding in the efficient allocation of resources and the implementation of tailored measures to control viral spread. Moreover, a comprehensive understanding of virus burden can contribute to the development of more effective vaccines and therapeutics, guiding research towards areas of greatest need. Monitoring viral burden enables timely detection of emerging pathogens and their variants, supporting rapid responses to prevent outbreaks and reduce their impact on communities. At the same time, PhIP technologies have strong potential to deepen our understanding of specific immune responses by identifying antibody fractions and targeted epitopes whose biological significance might otherwise be overlooked or underestimated. We believe that leveraging epitope libraries within PhIP platforms can substantially support global efforts to control viruses in human populations on a solid molecular basis.

## 4 Conclusions

1. Despite recognizing only a small fraction of potential viral epitopes, human IgG responses still may achieve broad coverage of virus species, genera, and families across the population (Figure 7).
2. Recognition of different epitopes of the same virus appears to vary considerably between individuals, and serological profiles may be highly individual.
3. Ability of an epitope to induce strong (high magnitude) response has only a limited effect on how often this epitope is recognized across the population. Also, high viral burden correlates with the frequency of immunogenic epitope recognition, but not with the strength of that recognition.
4. Serological profiling facilitates serological comparisons between populations.
5. We propose the **burden score** that helps normalize for differences in viral proteome representation in reference databases when evaluating immune burden.

**Figure 7.**
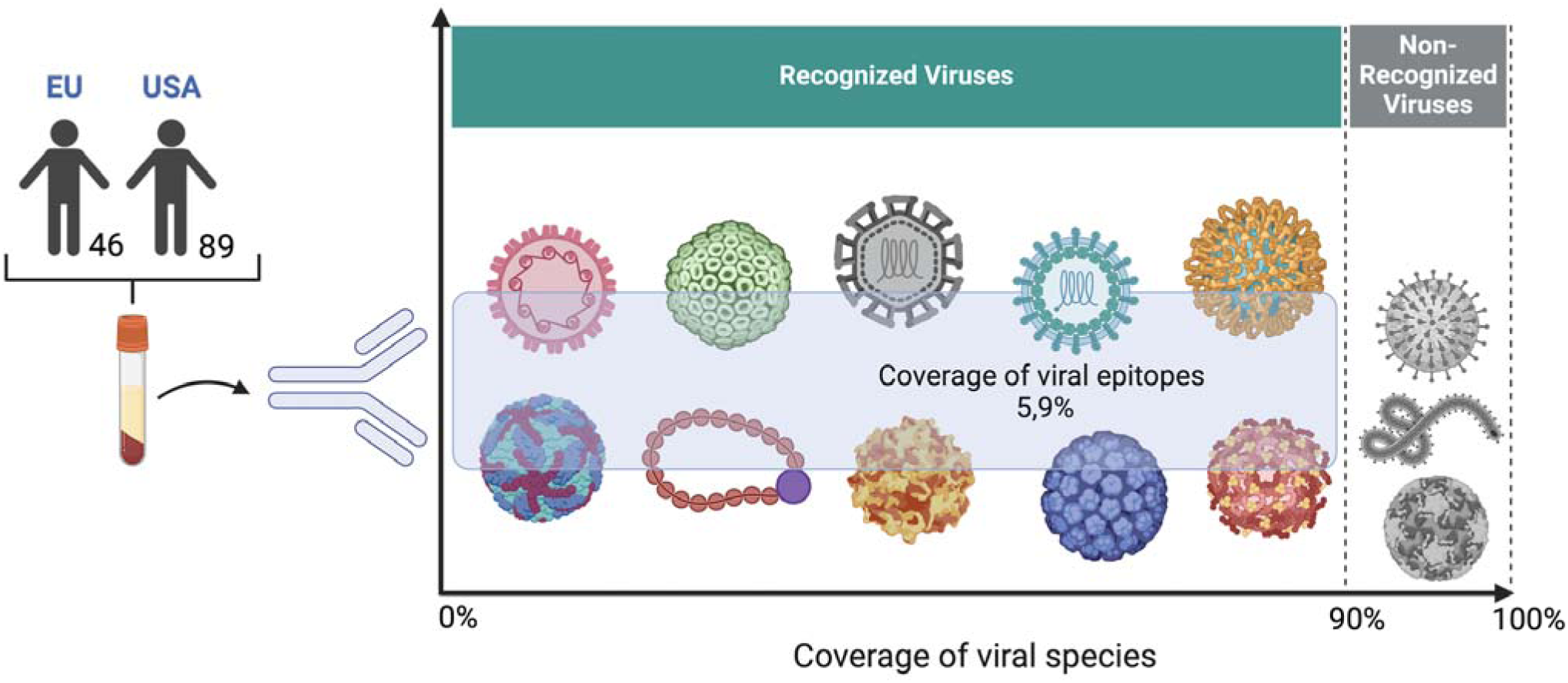
Human IgGs recognize 5.9% of viral epitopes but responses still achieve over 90% coverage of virus species, genera, and families across the population.

**Figure 8.**
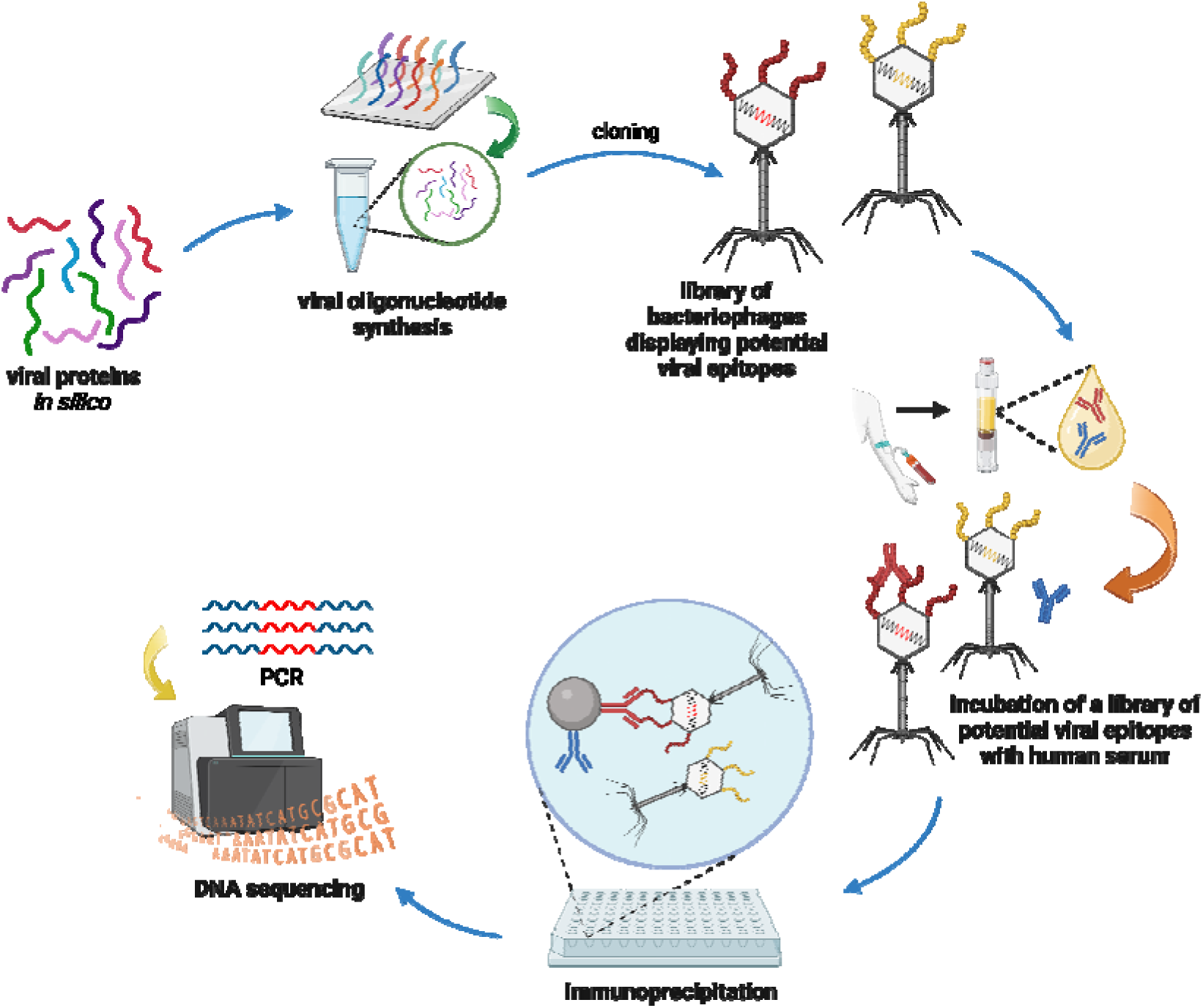
Overview of the Phage Display ImmunoPrecipitation technology used in presented research (modified from Xu et al.) [14]. Firstly (upper left) sequences of peptides representing viral proteome were downloaded, translated back to DNA code, synthetized and the library of bacteriophages displaying potential viral epitopes was created. Next the library was incubated with human sera creating bacteriophage-protein-IgG complexes that were immunoprecipitated. At this step viral proteins that were not recognized by IgG in human sera were washed out. Remaining phages were isolated and genome fragments coding presented viral epitope were sequenced and counted.

## 6 Materials & Methods

### 6.1 Library preparation

In June 2018, we conducted a search for viral proteins in UniProt (host: human, reviewed:yes, UniRef90) and subsequently downloaded protein sequences. From this dataset, 3,195 representatives of viral proteomes were fragmented into 56 amino acid long oligopeptides with 28 amino acid overlaps spanning the entire protein sequence. These sequences were then reverse-translated into oligonucleotide sequences using the most common *E. coli*-expressed codons. Stop codons were added, along with primers containing *EcoRI* and *HindIII* restriction sites[28]. The oligonucleotide sequences were synthesized (Agilent), and the resulting library was used to create a T7 phage library using the commercially available T7Select 415 Cloning kit (Novagen), following the manufacturer’s instructions. At this stage it was ensured that at least triple the number of synthesized sequences (over 150 000) was packed into phage capsid at the packaging stage.

### 6.2 Library purification

The T7 phage library expressing viral oligopeptides was cultured in LB broth at 37°C with shaking at 400 rpm. When bacterial clearing was observed, the bacterial culture was centrifuged at 8000 x g for 5 min, followed by sterile filtration through 0.22 μm cellulose acetate filters with a 3 μm glass fiber prefilter. NaCl was added to the filtrate up to a concentration of 0.5 M, along with PEG-8000 to a final concentration of 10% (v/v). The samples were then incubated at 0°C for 18 hours without shaking. After incubation, the samples were centrifuged at 15,000 x g for 15 minutes at 0°C. The supernatant was discarded, and the pellet was resuspended in Phage Extraction Buffer (20 mM Tris-HCl, 100 mM NaCl, 6 mM MgSO_4_, pH 8.0) at 0.01 times the volume of the initial sample for three hours. Phage counts were enumerated using the plating method.

### 6.3 Phage Immunoprecipitation (PhIP) sequencing

PhIP was conducted following published detailed procedures[12]. Briefly, 2 µl of each human serum sample was combined with a pre-prepared T7 phage library containing viral oligopeptides at a concentration of 10^10^ PFU/mL (equivalent to 2×10^5^ representations of each clone in the library) in a 96-well plate, suspended in Phage Extraction Buffer. The mixture was incubated overnight at 4°C with shaking at 400 rpm. After incubation, 20 µl of Dynabeads Protein A and Dynabeads Protein G (ThermoFisher) were added. The samples were then incubated for three hours at 4°C with shaking at 400 rpm. Following this incubation, the Dynabeads were separated using a magnetic rack and washed three times with Phage Wash Buffer (50 mM Tris-HCl, pH 7.5, 150 mM NaCl, 0.1% Tween-20). At this stage, phages displaying oligopeptides with epitopes recognized by IgG antibodies in the human serum were bound to the Dynabeads, while unbound IgG antibodies and phages with non-recognized oligopeptides were washed away. All samples and necessary controls were then suspended in water. After resuspension, all samples, controls, and input samples were subjected to two rounds of PCR. The first round used primers that replicated region of T7 genomes encoding a sequence of oligopeptide presented on the phage capsid and sequences complementary to 3’ ends of IDT for Illumina DNA/RNA UD Indexes (Illumina) which were used in the second round of PCR resulting in ready for sequencing amplicon library. Finally, all samples, controls, and input samples were pooled and sequenced using the NextSeq 550 System (Illumina, Figure 8).

### 6.4 NGS data processing

Data processing was conducted based on the original works of Xu et al[13,14], following a procedure closely aligned with the approach detailed in our previously published work[29]. Paired reads underwent de-multiplexing, merging, and removal of index sequences. The resulting amplicon sequences were then mapped to the original nucleotide library sequences using Bowtie2 software, utilizing the full list of oligonucleotides designed for library synthesis as indexes. The mapping was performed in end-to-end mode with specific options (“-q - 5 9 --no-unal --no-hd --no-sq --ignore-quals --mp 3 --rdg 150,100 --rfg 150,100 --score-min L,-0.6,-0.6”). The number of hits mapped to each reference sequence was counted, considering only the highest score for each read. The signal in each sample was calculated/normalized according to Formula (1):

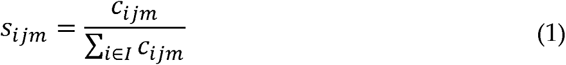

s - a signal of the *i-th* sequence in the *j-th* serum sample and *m-th* technical replicate
c - count, the number of reads mapped to the *i-th* sequence in the *m-th* technical replicate of the *j-th* serum sample
I - set of all reference sequences (used as indexes in mapping by the Bowtie2 software)

Input samples (n = 10, input controls) comprising amplified, purified, and sequenced phage libraries before immunoprecipitation, were used to evaluate background signals. For each sequence in each input sample, the log_10_ of the signal was calculated, along with the mean and standard deviation (SD) of the log10 signal across all input samples. These mean and SD values were then used for subsequent p-value calculations. To mitigate potential false positive results, the signal of undetected oligonucleotides in input sample was set to the minimum signal measured for that sample (count=1) and SD in oligopeptides that were recognized by low count (0 or 1) was set to average SD value. Oligopeptide sequences that were synthesized but not detected in any sample were excluded from further analysis. Full list of sequences with mean values in input group, SD and whether they were detected or not is in Supplementary Table 15.

For the tested samples, the log_10_(signal) was determined for each detected sequence (count > 0). To mitigate potential false positive results, the signal of undetected oligonucleotides in any sample was set to the minimum signal measured for that sample (count=1). Each log_10_(signal) in the tested samples was compared to the mean signal and SD of the corresponding sequence in the input samples, and the probability of a measured value or larger by chance was calculated using the R programming language (p-value, normal distribution). P-values were then adjusted for multiple hypotheses using the FDR method, separately for each tested sample[30]. Immunogenic (positive, significantly enriched) oligopeptides were defined as those with a non-zero count and an adjusted p-value < 0.05 in both technical replicates of a sample.Enrichment (relative signal) was calculated as the average signal of a particular sequence from both the technical replicates, divided by the mean signal in input samples of the same sequence, resulting in a signal ratio (relative signal). A no-serum negative control was included to identify nonspecific phage pull-downs that could lead to false-positive classification of immunogenic oligopeptides. An aliquot of the initial phage library was subjected to the complete PhIP procedure in parallel with the tested samples, with neutral buffer replacing serum. Oligopeptide abundance in this control was evaluated against the mean and standard deviation in the input-library samples using the same enrichment criteria as for the serum-containing samples.

We found 114 oligopeptides showing statistically significant increase in enrichment in comparison to input samples (adj. p < 0.05). However, only 2 of these oligopeptides were detected as immunogenic in presented results. The first one ‘UniRef90_Q1PDC7_oligopeptide_10’ was detected as immunogenic in 4 serum samples (2.98% of all samples) and ‘UniRef90_P37800_oligopeptide_6’ was detected as immunogenic in only 1 tested sample (0.75% of all samples). Full list of oligopeptides detected in negative control, results of its comparison to input samples along with means and SD are in Supplementary Table 16. These results were left in the presented analysis due to oligopeptides being recognized in less than 5% of samples tested. Additionally, we checked the phage concentration in wash fraction and original library after PhIP with dilution method in three randomly chosen samples and a control. Based on phage concentrations on three wash steps, we determined that there is less than 10% of phage contaminants in final elution of PhIP.

Importantly, oligonucleotides originally synthesized for the creation of the T7 phage library but not detected in any control or experimental samples were assumed either to be lost due to technical reasons or not enriched in any of the samples enough to be detected at used sequence depth. The overall charge, amino acid frequency, oligopeptide molecular mass and protein/species of origin of not detected oligopeptides were analyzed and compared to whole library. We present full list of sequences in Supplementary Table 1 and Supplementary Table 2.

### 6.5 Random control for comparison to randomly chosen oligopeptides

Control samples simulating the random selection of immunogenic oligopeptides from the original library underwent a two-step process. First, a data table was created that associated serum samples with the quantities of immunogenic oligopeptides detected in them. Next, oligopeptides were randomly selected from the original library in quantities corresponding to those determined in the first step. This process generated a dataset of 135 random control samples, each containing an average of 90 ± 21 oligopeptide sequences from the library set (Supplementary Data 1).

For the analysis of proteins, species, genera, and families of origin, the same assumption was applied as in the experimental samples: if an oligopeptide was recognized as immunogenic (significantly enriched, positive) in at least one serum sample, the corresponding viral protein, species, genus, and family of origin for that oligopeptide were marked as “recognized” and “immunogenic” in that serum sample.

### 6.6 Random control for comparison to randomly chosen immunogenic oligopeptides

Control samples simulating the random selection of immunogenic oligopeptides from the list of experimentally detected and significantly enriched ones were generated in three steps. First, a data table was created to show the quantities of immunogenic oligopeptides detected in specific serum samples. Next, oligopeptides were randomly chosen from a list of unique, experimentally detected immunogenic oligopeptides, in quantities matching those in the first step. This process established a dataset of 135 random control samples, each containing an average of 90 ± 21 randomly chosen oligopeptides from the set of experimentally detected immunogenic ones. In the third step, the *in silico* simulation from the second step was repeated 99 additional times, for a total of 100 simulations. For the analysis of proteins, species, genera, and families of origin, the same assumption was applied as in section 6.5 of the Materials & Methods: if an oligopeptide was recognized as immunogenic (significantly enriched, positive) in at least one serum sample, the corresponding viral protein, species, genus, and family of origin for that oligopeptide were marked as “recognized” and “immunogenic” in that serum sample.

### 6.7 Recognition viral proteins, species genera and families of origin

Protein codes for the downloaded proteins used in library creation were obtained following the procedure outlined in section 6.1 of the Materials & Methods. Species designations were obtained from the UniRef90 database, specifically from the “Organism” field at the time of the initial library preparation. It is important to note that certain proteins represent specific viral subtypes. To prevent false negatives, we merged descriptors of virus species if the species were the same but the viral subtypes were differed (see Supplementary Table 17). Viral genera and families were identified using taxonomy information available on the UniProt online resource. Virus names were imported directly from the UniProt “Organism” field (for instance, rhinovirus entries appeared in our study under Enterovirus-related designations). However, it is important to acknowledge that our species descriptions may not fully align with the current UniRef90 and taxonomy resources at the time of publication. We provide the original descriptions so that readers can download them and extract results of interest, adapting their analysis to their needs and the current taxonomy (Supplementary Table 4).

### 6.8 Shannon Diversity Index

The Shannon Diversity Index (SDI) was calculated using the originally published equation[31]. Each proportion was determined by dividing the total number of times each entity (oligopeptide, protein, species, genus, or family) was recognized as immunogenic across all samples by the sum of all entities recognized as immunogenic across all samples. The resulting SDI was calculated for the experimentally measured samples and for 100 sets of randomly selected control samples from immunogenic oligopeptides (as described in section 6.6 of the Materials & Methods). P-values were calculated by comparing the index values of the measured experimental samples with the means and standard deviations from the 100 sets of control samples, using a normal distribution and a two-tailed test. Exact p-values for oligopeptides and proteins were reported by the R programming language as below 2×10^-16^. Exact p-values for virus species, genera, and families were also reported.

### 6.9 Analysis of over- and underrepresented oligopeptides, proteins, species, genera, and families

We conducted the analysis of over- and underrepresentation in three-step process. First, we used the simulated *in silico* samples described in section 6.6 of the Materials & Methods. Next, we assessed the probability that the observed fraction of serum samples recognizing each immunogenic oligopeptide was due to chance, using a normal distribution and a two-sided test. The resulting p-values were adjusted for multiple hypotheses using the FDR method[30]. Oligopeptides with adjusted p-values < 0.05 were considered significantly overrepresented or underrepresented. Similar analyses were conducted for proteins, species, genera, and families with the same assumption applied as in the experimental samples: if an oligopeptide was recognized as significantly enriched (immunogenic) in at least one serum, the viral protein, species, genus, and family of origin for that oligopeptide were marked as “recognized” in that serum sample. The calculated values are presented in Supplementary Tables 5-9.

### 6.10 Analysis of differences between geographically distinct groups

To assess overrepresentation and underrepresentation among geographically distinct groups, we counted the number of serum samples recognizing an oligopeptide as immunogenic for each group (Supplementary Table 12). In order to compare sources of variation in our data, we compared deviance in null binomial model without splitting between groups for total deviance and then with splitting by US and PL groups for in-group deviance. For evaluation of oligopeptides and virus species significantly varying between groups, for each recognized immunogenic oligopeptide, we calculated the p-value between these counts using a two-sided binomial distribution test. Due to the absence of a suitable control group representing the general population at the time of writing, we conducted the p-value calculation between the US group and the PL group, and vice versa, to address the non-symmetrical binomial distribution model and minimize false positives. The higher p-value from these comparisons was selected for analysis. P-values were then adjusted for multiple hypotheses using the FDR method[30]. Adjusted p-values below 0.05 indicated oligopeptides overrepresented in the respective group. The species-level analysis followed the same protocol, considering the number of serum samples recognizing a virus species as immunogenic for each group (Supplementary Table 13).

### 6.11 Cross-reaction analysis for burden score calculation

The normalization addresses the problem of non-equal representation of viral species in original database. Viral species that are heavily researched and analyzed are represented by many proteomes in databases have higher chance of cross-reactions than others, as seen by correlation.

Our process assumes that if cross-reactions are primarily responsible for detecting most IgG-oligopeptide interactions, then the likelihood of detecting an oligopeptide representing a specific virus species is similar to randomly selecting an oligopeptide representing that virus species from the original input library. Burden score was determined by comparing the fraction of all recognized immunogenic oligopeptides representing a specific virus species,, to the fraction of the original input library represented by that same virus species. This comparison is mathematically presented in Formula (2).

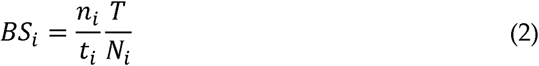

BS_i_ – burden score of *i-th* virus species
n_i_ – the number of oligopeptides representing the *i-th* virus species in all tested sera
t – total number of immunogenic oligopeptides identified in all sera
N_i_ – the number of oligopeptides representing the *i-th* virus species in original library of oligopeptides downloaded from UniProt
T – total number of oligopeptides representing all virus species in the original library of oligopeptides downloaded from UniProt

For calculations of p-value we used binomial model with probability equal to ratio of the number of sequences representing i-th virus species to all sequences in original library (*N_i_/T*). The sum of all detected sequences representing i-th virus species and sum of all detected sequences in the presented research were used in binomial model as the number of tested events and total number of events. P-values calculated by SciPy package and Python Programming Language[32]. P-values adjusted for multiple hypotheses with were calculated using Benjamini-Hochberg method (false discovery rate) in statsmodel package (‘fdr_bh’) [30,33].

Like Xu et al. 2015, we calculated the virus specific threshold (cut-off) by calculating linear regression between log_10_(average signal) and log_10_(number of unique proteins in library) for each virus species. Using calculated slope and threshold value of three for herpes simplex virus 1, like Xu et al., we calculated for each virus species a threshold value[14]. Min. value of two was set to avoid spurious hits. If we detected in any sample equal or more immunogenic oligopeptides from a given virus species than the virus specific threshold for this species, this serum sample was marked as exposed to a virus species in the past.

### 6.12 Calculations and correlations

All calculations were performed, unless stated explicitly otherwise, using the R programming language and the dplyr package, unless otherwise specified in the Materials & Methods[34,35]. GraphPad Software (Boston, Massachusetts, USA) was extensively used throughout our research for data evaluation and plot creation[36]. Correlations are presented in the format: Pearson’s r(x)=y, 95% CI z to w, p=k, spearman’s ρ(x) = l, p=k. In this format, “r” represents Pearson’s correlation coefficient, “x” denotes the degrees of freedom, “y” is the value of Perason’s r, “z” and “w” are the lower and upper boundaries of the 95% confidence interval (CI), respectively, “p” indicates the p-value, “k” is the numerical value of p-value, “ρ” is Spearman’s correlation coefficient, “l” is the value of ρ.

### 6.13 Blood samples

Blood serum was obtained from 46 healthy adult volunteers from both Poland from 89 individuals from the US, the latter ones commercially sourced with Prof. D. C. Nelson (IBBR, University of Maryland, MD, USA), the US blood samples were collected in a clotting system.

Polish cohort blood samples were collected in test tubes (BD SST II Advance), left to clot for 1 hour at room temperature (RT), and separated from the clot by centrifugation (15 min, 2000 g, RT) and then stored at – 20°C for further use.

### 6.14 Demographic information on the participants is available in Supplementary Table 11); age data are available only for the US cohort while PL cohort had been ultimately anonymized without age data (that are considered an indirect identifier), which limits the possibility of analyzing age-associated effects in this study. Bioethics statements

The study was conducted in accordance with the principles of the Declaration of Helsinki. The research was approved by the local Bioethical Commission of the Regional Specialist Hospital in Wroclaw (approval number: KB/02/2017). Blood collection in During the Regional Specialist Hospital in Wroclaw was conducted with individual patient interview, all information about the study was provided and written consent was obtained from each participant. The written consent form was accepted by the local Bioethical Commission of the Regional Specialist Hospital in Wroclaw (approval number: KB/02/2017).

### 6.15 Data availability

Original data from sequencing in fastq format used in this manuscript is accessible in European Nucleotide Archive under acc no.: PRJEB81945 (project id) or ERP165715 (study id).

## 7 Acknowledgments

## 7.1 Financial statement

This work was supported by the National Science Centre in Poland: grant no. 2019/35/B/NZ7/01824.

## 7.3 Author contributions

Marek Adam Harhala – conceptualization, data curation, formal analysis, investigation, methodology, software, supervision, visualization, writing – original draft, writing – review & editing

Katarzyna Gembara - data curation, investigation, methodology, software, supervision, validation, visualization

Daniel C. Nelson -data curation, validation, visualisation, writing – review & editing

Andrzej Konieczny - investigation, resources,

Natalia Jędruchniewicz - investigation, resources, project administration,

Izabela Rybicka - data curation, investigation, methodology,

Krystyna Dąbrowska - conceptualization, data curation, funding acquisition, investigation, methodology, project administration, resources, supervision, validation, visualization, writing – review & editing

## 10 Supplementary Data

**Supplementary Data 1:** This file contains all oligopeptides tested in the study, with their names and sequences provided in the fasta format.

**Supplementary Table 1:** This table lists all immunogenic oligopeptides identified in the study. An oligopeptide is considered immunogenic if it is recognized by IgG in at least one of tested serum samples, meaning that the signal in at least one sample is significantly higher than the input samples (libraries before immunoprecipitation) in both technical replicates. The column “oligopeptide.name” represents the names of the oligopeptides as used in our research. The column “protein.code.UniRef90” provides the protein code as downloaded from the UniRef90 resource during library creation. The “oligopeptide.number” column indicates the starting position of the 56 amino acid oligopeptide in the protein sequence, with each oligopeptide beginning every 28 amino acids, starting from the first one. The column “n” shows the number of serum samples that recognized a specific oligopeptide as immunogenic. Finally, the “ratio” column displays the percentage of serum samples that recognized a specific oligopeptide as immunogenic.

**Supplementary Table 2:** This table presents counts of immunogenic (recognized) oligopeptides identified in this study. Column ‘olgnname’ is sequence name, ‘mean.control.log.signal’, ‘control.sd.log.signal’ list log_10_(signal) and sd in controls before immunoprecipitation, ‘count’, ‘log.signal’, ‘relative.signal’ and ‘p.value’ list the number of alignments, log_10_(signal), relative signal and p-value of a measurement in a respective sample in first (x) and second (y) technical replicate. Columns ‘sum.x’ and ‘sum.y’ list the number of all aligned sequences in a respective sample in first and second technical replicate, respectively. Columns ‘p.value’ an ‘adj.p’ show p-value and p-value adjusted for multiple hypotheses (fdr=5%) of a measurement after accounting for both technical replicates. Column ‘sample’ lists the numerical designation of a tested sample.

**Supplementary Table 3:** This table presents the expected fractions and quantities of viral oligopeptides, proteins, species, genera, and families recognized as “immunogenic” when detected randomly from the list of all oligopeptides in the input samples, provided in a .csv format. Each row represents one iteration of a random control sample; for details, see section 6.5 in Materials & Methods. The “X.ratio” columns represent the fractions of all tested X (where X could be oligopeptides, proteins, species, genera, or families) that would be marked as immunogenic when detected randomly. The “n.X” columns indicate the quantities of X that would be marked as immunogenic under random detection. P-values for differences between the mean and standard deviation (SD) of the simulated and measured values, as presented in Figure 2A, are as follows: p<2×10^-16^ (oligopeptides), p<2×10^-16^ (proteins), p<1×10^-6^ (species), p<0.01 (genera), and p<0.01 (families).

**Supplementary Table 4:** This table presents the complete list of oligopeptides used in the research, including their names (“oligopeptide.name”), protein codes (“protein.code”), descriptions (“description”), species (“species”), genera (“genus”), subfamilies (“subfamily”, if applicable), and families (“family”) of origin. The data were collected at the time of library creation, as listed in UniRef90 resource.

**Supplementary Table 5:** This table lists all viral oligopeptides recognized as immunogenic. The column “oligopeptide.name” contains the names of the oligopeptides as used in our research. The columns “n” and “ratio” represent the number and fraction of serum samples, respectively, that recognized each specific oligopeptide as immunogenic. Oligopeptides not detected as immunogenic (n=0) are not included. The columns “mean.control” and “sd.control” show the mean and SD, respectively, of the expected (simulated *in silico*) fractions of serum samples that recognized each specific oligopeptide as immunogenic. The “p.values” column presents the p-values calculated between the expected (“mean.control” and “sd.control”) and observed values using a normal distribution and two-tailed test. The “adj.p” column shows p-values adjusted for multiple hypotheses using the FDR method. The “flag” column indicates whether the immunogenic oligopeptides were overrepresented, underrepresented (adj.p<0.05) or normally represented (adj. p>0.05) in comparison to the simulated set of controls (see section 6.9 in Materials & Methods).

**Supplementary Table 6:** This table lists all viral proteins recognized as immunogenic. A viral protein was marked as immunogenic if any oligopeptide within that protein was detected as immunogenic in at least one of tested sera samples. The column “protein.code” represents protein code as listed in the UniRef90 resource, which was used in our research. The columns “n” and “ratio” indicate the number and fraction of serum samples, respectively, that recognized each specific protein as immunogenic. Proteins not detected as immunogenic (n=0) are not included. The columns “mean.control” and “sd.control” show the mean and SD, respectively, of the expected (simulated *in silico*) fractions of serum samples that recognized each specific oligopeptide as immunogenic. The “p.values” column presents the p-values calculated between the expected (“mean.control” and “sd.control”) and observed values using a normal distribution and two-tailed test. The “adj.p” column shows p-values adjusted for multiple hypotheses using the FDR method. The “flag” column indicates whether the immunogenic oligopeptides were overrepresented, underrepresented (adj.p<0.05) or normally represented (adj. p>0.05) in comparison to the simulated set of controls (see section 6.9 in Materials & Methods).

**Supplementary Table 7:** This table lists all virus species recognized as immunogenic. A virus species was marked as immunogenic if any oligopeptide belonging to that species was detected as immunogenic in at least one of tested sera samples. The column “species” represents the virus species as listed in the UniRef90 resource (in the “organism” section) at the time of library creation for each protein. The columns “n” and “ratio” indicate the number and fraction of serum samples, respectively, that recognized each specific virus species as immunogenic. Virus species not detected as immunogenic (n=0) are not included. The columns “mean.control” and “sd.control” show the mean and SD, respectively, of the expected (simulated *in silico*) fractions of serum samples that recognized each specific oligopeptide as immunogenic. The “p.values” column presents the p-values calculated between the expected (“mean.control” and “sd.control”) and observed values using a normal distribution and two-tailed test. The “adj.p” column shows p-values adjusted for multiple hypotheses using the FDR method. The “flag” column indicates whether the immunogenic oligopeptides were overrepresented, underrepresented (adj.p<0.05) or normally represented (adj. p>0.05) in comparison to the simulated set of controls (see section 6.9 in Materials & Methods).

**Supplementary Table 8:** This table lists all viral genera recognized as immunogenic. A genus was marked as immunogenic if any oligopeptide belonging to that genus was detected as immunogenic in at least one of tested serum samples. The column “genus” lists the genera representing the tested viral proteins, as recorded in the taxonomy resource of the UniProt database at the time of library creation. The columns “n” and “ratio” represents the amount and the fraction of serum samples, respectively, that recognized each specific genus as immunogenic. Genera not detected as immunogenic (n=0) are not included. The columns “mean.control” and “sd.control” show the mean and SD, respectively, of the expected (simulated *in silico*) fractions of serum samples that recognized each specific oligopeptide as immunogenic. The “p.values” column presents the p-values calculated between the expected (“mean.control” and “sd.control”) and observed values using a normal distribution and two-tailed test. The “adj.p” column shows p-values adjusted for multiple hypotheses using the FDR method. The “flag” column indicates whether the immunogenic oligopeptides were overrepresented, underrepresented (adj.p<0.05) or normally represented (adj. p>0.05) in comparison to the simulated set of controls (see section 6.9 in Materials & Methods).

**Supplementary Table 9:** This table lists all viral families recognized as immunogenic. A family was marked as immunogenic if any oligopeptide belonging to that family was detected as immunogenic in at least one of tested serum samples. The column “family” lists the viral families representing the tested viral proteins, as recorded in the taxonomy resource of the UniProt database at the time of library creation. The columns “n” and “ratio” represents the number and fraction of serum samples, respectively, that recognized each specific family as immunogenic. Families not detected as immunogenic (n=0) are not included. The columns “mean.control” and “sd.control” show the mean and SD, respectively, of the expected (simulated *in silico*) fractions of serum samples that recognized each specific oligopeptide as immunogenic. The “p.values” column presents the p-values calculated between the expected (“mean.control” and “sd.control”) and observed values using a normal distribution and two-tailed test. The “adj.p” column shows p-values adjusted for multiple hypotheses using the FDR method. The “flag” column indicates whether the immunogenic oligopeptides were overrepresented, underrepresented (adj.p<0.05) or normally represented (adj. p>0.05) in comparison to the simulated set of controls (see section 6.9 in Materials & Methods).

**Supplementary Table 10:** This table lists sequences with the maximum relative signal for a given immunogenic oligopeptide from all signals detected in tested samples. Columns ‘oligopeptide.name’ lists the name of a sequence, ‘max.relative.signal’ lists the highest signal from all samples for a respective oligopeptide. ‘number.of.positive.samples’ and ‘ratio’ shows the number of samples and the fraction of all samples that detected the respective oligopeptide as immunogenic.

**Supplementary Table 11:** This table lists reported composition of tested groups in section 2.4. Data and samples on USA group was obtained from BioIVT company.

**Supplementary Table 12:** This table details the geographical disparity in unique and shared immunogenic oligopeptides. It lists all oligopeptides recognized as immunogenic in serum samples from the US group (n=88) and the Polish (PL) group (n=46). The column “oligopeptide.name” represent the name of each oligopeptide as used in our research, The “n.X” columns show the number of serum samples in the X group that recognized a specific oligopeptide as immunogenic, while the “fraction.X” columns indicate the fraction of serum samples in the X group that recognized the oligopeptides as immunogenic. The “p.value” column provides the exact p-value (binomial distribution, two-tailed test) representing the probability that the observed difference between the groups is accidental. The “adj.p” column shows p-values adjusted for multiple hypotheses using the FDR method. The “overrepresented” column indicates the group in which a specific oligopeptide is overrepresented, or “NO” if the adjusted p-value is greater than 0.05. The “group” columns flag the groups that recognized a specific oligopeptide as immunogenic, with “X.only” indicating oligopeptides unique to the X group and “BOTH” indicating oligopeptides detected in both groups. The table is sorted by increasing values in the “adj.p” column.

**Supplementary Table 13:** Geographical disparity in unique and shared recognized viruses. This table lists all virus species recognized in serum samples from the US group (n=88) and the Polish (PL) group (n=46). The column labeled “species” indicates the name of each virus species used in our research. The “n.X” columns represent the number of serum samples that recognized a specific species as immunogenic in group X, while the “fraction.X” columns represent the fraction of serum samples that recognized the specific species as immunogenic in group X. The “p.value” column provides the exact p-value (binomial distribution, two-tailed test) comparing the PL and US groups. The “adj.p” column shows the p-values adjusted for multiple hypotheses testing using the FDR method. The “overrepresented” column indicates a group in which the specific virus species is overrepresented, or “NO” if the adjusted p-value >0.05. Finally, the “group” columns use flags to show which groups recognized a specific virus species as immunogenic (“X.only” for oligopeptides unique for group X, and “BOTH” for oligopeptides detected in both groups). The table is sorted by ascending values in the “adj.p” column.

**Supplementary Table 14:** The burden of virus species impacting the human population – development of a normalization approach accounting for oligopeptide frequency in the tested library. This table presents a list of virus species that impose a burden on the immune system, normalized to account for the frequency of oligopeptides in the tested library. The column “n.library” indicates the number of oligopeptides representing each specific virus in the original library. The column “fraction.library” shows the fraction of oligopeptides representing each specific virus in the original library. The “fraction.results” column indicates the fraction of oligopeptides representing each virus in the list of all recognized immunogenic oligopeptides across all serum samples. The “burden.score” column displays the odds ratio of recognizing oligopeptides representing a specific virus species as immunogenic as compared to finding them in original library by chance (with an burden score value of 1 indicating odds of 1:1). The “n.sera.samples” column represents the number of serum samples that recognized a specific virus as immunogenic, based on the criterion that at least three oligopeptides representing a virus species must be recognized in a serum sample for it to be considered immunogenic. The “fraction.exposed” column indicates the fraction of serum samples that recognized a virus species as immunogenic. The “p.value” column presents the exact p-values (binomial distribution, two-sided) calculated to assess whether the observed burden score is statistically significant. The “adj.p” column shows the p-value adjusted for multiple hypothesis testing using the FDR method.

**Supplementary Table 15:** This table lists all sequences tested. Names are in ‘oligopeptide.name’ column, negative log10 of mean signal in control samples and its SD is in ‘mean.control.log.signal’ and “control.sd.log.signal”, respectively. Fraction of control samples in which specific oligopeptide was detected is in ‘ratio.detected’ column and ‘flag’ column lists if respective oligopeptide was detected in any of tested samples in any run.

**Supplementary Table 16:** This table lists virus oligopeptides detected in true negative controls as described in Section 6.4. Column ‘oligopeptide.name’ lists sequence names, “mean.control.log.signal” and “control.sd.log.signal” lists log10(signal) values of respective sequences in the control samples, samples prior to precipitation that are reference point for measuremence of statistical significance. Columns “count” and “log.neg_control.signal” show the number of reads in the true negative sample determined to come from a respective sequence and its respective log10(signal). “P-value” and “adjusted_p” columns show probability of accidental signal of such strength in comparison to controls as calculated by binomial model. “adjusted_p” shows p-values adjusted for multiple hypotheses (fdr=5%). Details are described in Section 6.4.

**Supplementary Table 17:** This table lists virus species that were originally classified with subtypes but were merged for the purposes of this study (see section 6.7 of Materials & Methods for details). The “original.species” column represents the name of the virus species as listed in the “organism” field for a protein representative in the UniRef90 resource of the UniProt database. The “species” column shows the species descriptor used in this research.

## References

1. Liston A, Humblet-Baron S, Duffy D, Goris A. Human immune diversity: from evolution to modernity. Nature Immunology. Nature Research; 2021. p. 1479–89. doi:10.1038/s41590-021-01058-1 PubMed PMID: 34795445.

2. Quyen TL, Ngo TA, Bang DD, Madsen M, Wolff A. Classification of Multiple DNA Dyes Based on Inhibition Effects on Real-Time Loop-Mediated Isothermal Amplification (LAMP): Prospect for Point of Care Setting. Front Microbiol. 2019;10(October):1–12. doi:10.3389/fmicb.2019.02234

3. Yitbarek K, Abraham G, Girma T, Tilahun T, Woldie M. The effect of Bacillus Calmette–Guérin (BCG) vaccination in preventing sever infectious respiratory diseases other than TB: Implications for the COVID-19 pandemic. Vaccine. 2020;38(41):6374–80. doi:10.1016/j.vaccine.2020.08.018 PubMed PMID: 32798142.

4. Sakuraba A, Haider H, Sato T. Population difference in allele frequency of hla-c*05 and its correlation with covid-19 mortality. Viruses. 2020;12(11). doi:10.3390/v12111333 PubMed PMID: 33233780.

5. Shi P, Dong Y, Yan H, Zhao C, Li X, Liu W, et al. Impact of temperature on the dynamics of the COVID-19 outbreak in China. Science of the Total Environment. 2020;728(77):138890. doi:10.1016/j.scitotenv.2020.138890 PubMed PMID: 32339844.

6. Nickbakhsh S, Mair C, Matthews L, Reeve R, Johnson PCD, Thorburn F, et al. Virus-virus interactions impact the population dynamics of influenza and the common cold. Proc Natl Acad Sci U S A. 2019;116(52):27142–50. doi:10.1073/pnas.1911083116 PubMed PMID: 31843887.

7. Vidarsson G, Dekkers G, Rispens T. IgG subclasses and allotypes: From structure to effector functions. Front Immunol. 2014;5(OCT):1–17. doi:10.3389/fimmu.2014.00520 PubMed PMID: 25368619.

8. Fanning LJ, Connor AM, Wu GE. Development of the immunoglobulin repertoire. Clin Immunol Immunopathol. 1996;79(1):1–14. doi:10.1006/clin.1996.0044 PubMed PMID: 8612345.

9. Thomas DM, Sturdivant R, Dhurandhar N V., Debroy S, Clark N. A Primer on COVID-19 Mathematical Models. Obesity. 2020;28(8):1375–7. doi:10.1002/oby.22881 PubMed PMID: 32386464.

10. Younes AB, Hasan Z. COVID-19: Modeling, prediction, and control. Applied Sciences (Switzerland). 2020;10(11):1–14. doi:10.3390/app10113666

11. Mohan D, Wansley DL, Sie BM, Noon MS, Baer AN, Laserson U LHB. PhIP-Seq Characterization of Serum Antibodies Using Oligonucleotide Encoded Peptidomes. Nat Protoc. 2018;13(9):1958–78. doi:10.1038/s41596-018-0025-6

12. Larman HB, Laserson U, Querol L, Verhaeghen K, Solimini NL, Xu GJ, et al. PhIP-Seq characterization of autoantibodies from patients with multiple sclerosis, type 1 diabetes and rheumatoid arthritis. J Autoimmun. 2013;43:1–9. doi:10.1016/j.jaut.2013.01.013 PubMed PMID: 23497938.

13. Larman HB, Laserson U, Querol L, Verhaeghen K, Solimini NL, Xu GJ, et al. PhIP-Seq characterization of autoantibodies from patients with multiple sclerosis, type 1 diabetes and rheumatoid arthritis. J Autoimmun. 2013;43:1–9. doi:10.1016/j.jaut.2013.01.013 PubMed PMID: 23497938.

14. Xu GJ, Kula T, Xu Q, Li MZ, Vernon SD, Ndung’u T, et al. Comprehensive serological profiling of human populations using a synthetic human virome. Science (1979). 2015;348(6239):1–23. doi:10.1126/science.aaa0698 PubMed PMID: 26045439.

15. Shrock EL, Shrock CL, Elledge SJ. VirScan: High-throughput Profiling of Antiviral Antibody Epitopes. Bio Protoc. 2022;12(13). doi:10.21769/BioProtoc.4464

16. Shrewsbury JV, Vitus ES, Koziol AL, Nenarokova A, Jess T, Elmahdi R. Comprehensive phage display viral antibody profiling using VirScan: potential applications in chronic immune-mediated disease. J Virol. 2024;98(11). doi:10.1128/jvi.01102-24

17. Monaco DR, Kottapalli S V., Breitwieser FP, Anderson DE, Wijaya L, Tan K, et al. Deconvoluting virome-wide antibody epitope reactivity profiles. EBioMedicine. 2022;75. doi:10.1016/j.ebiom.2021.103747

18. Nowak L. Ludność. Stan i struktura demograficzno-społeczna. Główny Urząd Statystyczny; 2011.

19. World Health Organization. https://data.who.int/countries/616. 2025. World Health Organization 2025, Poland [Country overview].

20. World Health Organization. https://data.who.int/countries/840. 2025. World Health Organization 2025, United States of America [Country overview].

21. Global HIV statistics [Internet]. [cited 2024 Mar 22]. Available from: https://www.unaids.org/en/resources/fact-sheet

22. Lidenge SJ, Yalcin D, Bennett SJ, Ngalamika O, Kweyamba BB, Mwita CJ, et al. Viral Epitope Scanning Reveals Correlation between Seasonal HCoVs and SARS-CoV-2 Antibody Responses among Cancer and Non-Cancer Patients. Viruses. 2024;16(3). doi:10.3390/v16030448

23. Venkataraman T, Valencia C, Mangino M, Morgenlander W, Clipman SJ, Liechti T, et al. Analysis of antibody binding specificities in twin and SNP-genotyped cohorts reveals that antiviral antibody epitope selection is a heritable trait. Immunity. 2022;55(1). doi:10.1016/j.immuni.2021.12.004

24. Wang L, Candia J, Ma L, Zhao Y, Imberti L, Sottini A, et al. Serological responses to human virome define clinical outcomes of Italian patients infected with SARS-CoV-2. Int J Biol Sci. 2022;18(15). doi:10.7150/ijbs.78002

25. Panda S, Ding JL. Natural Antibodies Bridge Innate and Adaptive Immunity. The Journal of Immunology. 2014;194(1). doi:10.4049/jimmunol.1400844

26. Zarrineh M, Mashhadi IS, Farhadpour M, Ghassempour A. Mechanism of antibodies purification by protein A. Analytical Biochemistry. 2020. doi:10.1016/j.ab.2020.113909

27. Krishna C, Chowell D, Gönen M, Elhanati Y, Chan TA. Genetic and environmental determinants of human TCR repertoire diversity. Immunity and Ageing. 2020;17(1). doi:10.1186/s12979-020-00195-9

28. Maloy, S., V. Stewart and RT. Genetic analysis of pathogenic bacteria A laboratory manual. Cold Spring Harbor Laboratory Press; 1996. 603 p.

29. Harhala MA, Gembara K, Baniecki K, Pikies A, Nahorecki A, Jędruchniewicz N, et al. Experimental Identification of Cross-Reacting IgG Hotspots to Predict Existing Immunity Evasion of SARS-CoV-2 Variants by a New Biotechnological Application of Phage Display. Viruses. 2023 Dec 29;16(1):58. doi:10.3390/v16010058

30. Benjamini Y, Hochberg Y. Controlling the False Discovery Rate: A Practical and Powerful Approach to Multiple Testing. Journal of the Royal Statistical Society. 1995;57(1):289–300.

31. Shannon CE, Weaver W. THE MATHEMATICAL THEORY OF COMMUNICATION. 10th ed. University of Illinois Press; 1964.

32. Virtanen P, Gommers R, Oliphant TE, Haberland M, Reddy T, Cournapeau D, et al. SciPy 1.0: fundamental algorithms for scientific computing in Python. Nat Methods. 2020;17(3). doi:10.1038/s41592-019-0686-2

33. Seabold S, Perktold J. Statsmodels: Econometric and Statistical Modeling with Python. In: Proceedings of the 9th Python in Science Conference. 2010. doi:10.25080/majora-92bf1922-011

34. Wickham H, François R, Henry L, Müller K, Vaughan D. dplyr: A Grammar of Data Manipulation. https://CRAN.R-project.org/package=dplyr; 2023.

35. R Core Team. R: A Language and Environment for Statistical Computing [Internet]. Vienna: R Foundation for Statistical Computing; 2024 [cited 2025 Mar 6]. Available from: https://www.R-project.org/

36. GraphPad Software. GraphPad Prism for Windows. Boston: GraphPad software; 2024.

